# Neurological effects induced by micro- and nanoplastics in fish: a systematic review and meta-analysis

**DOI:** 10.1101/2025.06.07.658168

**Authors:** Querusche Klippel Zanona, Matheus Gallas-Lopes, Gabriel Alves Marconi, Isabela Nachtigall Lazzarotto, Angelo Piato, Ana Paula Herrmann, Maria Elisa Calcagnotto

## Abstract

Plastic is considered an inert material with high durability and minimal to virtually no decomposition. However, when released into the environment, they can degrade into very small particles, forming micro- and nanoplastic particles (MNP). This systematic review and meta-analysis synthesized the evidence from controlled preclinical studies to investigate the neurological effects of MNPs in fish. Following a pre-registered protocol, we searched PubMed, Web of Science, and Scopus for studies exposing fish to virgin MNPs under controlled conditions, reporting behavioral or neurochemical outcomes relevant to central nervous system function. Data were synthesized using a hierarchical random-effects model with robust variance estimation. Fifty-nine studies, comprising 723 comparisons across 13 behavioral and neurochemical outcomes, were included in the meta-analysis. The analysis accounted for correlated effect sizes within shared control groups; and nested identifiers for study, control group, and effect size. Results indicated high heterogeneity and no consistent effect of the MNP exposure on behavioral or neurochemical parameters, except for reduced traveled distance in sensory-motor assays. Meta-regression examined whether developmental stage, exposure duration, and MNP size or concentration moderate these effects. No significant moderators were identified, except for catalase activity, where longer exposure reduced enzyme activity in larvae. The findings should be interpreted with caution, as reporting quality was generally low, with key methodological details often omitted. Additionally, publication bias was found for several outcomes, and influential case analyses revealed that a few studies disproportionately affected the overall estimates. Further studies are needed to clarify the impact of MNPs on fish neurobiology and behavior.

## 1. Introduction

Plastic is considered an inert and highly durable material, qualities that have made it a valuable resource and led to its widespread replacement of glass bottles, paper packaging, wood, and metal products across various industries [1]. In 2022, approximately 400 million tonnes of plastics were produced globally [2]. Nearly two-thirds of all plastic produced is used in relatively short-lived products, such as packaging, consumer goods, and textiles [1]. This situation is concerning because plastics have an extremely low rate of biological decomposition [3,4], positioning plastic pollution among the greatest environmental challenges of the 21st century [5].

Waste management faces considerable difficulties, as only 9% of all plastic produced is effectively recycled. In 2022 alone, global plastic waste generation reached 285.1 million tonnes, with 31.6% incinerated, 13.3% recycled, 2.3% traded, and 48.2% entering the environment through landfill (36.2%), mismanagement (10.4%), or losses during production and transportation (6.2%) [2]. Under current policies, plastic leakage from sources such as littered plastic, synthetic textiles, vehicle tyres, and plastic pellets to terrestrial and aquatic environments is projected to increase by 50% between 2020 and 2040 [5]. These findings highlight the urgent need for effective policies and practices based on responsible consumption and production, as outlined in Sustainable Development Goal (SDG) 12, as the United Nations Environment Assembly resolution to end plastic pollution [6]. In addition, eliminating plastic pollution from aquatic environments contributes to SDG 6 (clean water and sanitation) and SDG 14 (life below water).

Microplastics are defined as plastic particles smaller than 5 mm, originating either from primary sources - intentionally manufactured small particles - or from secondary sources, fragmented from larger plastic debris [7,8]. Although initially detected in aquatic environments, microplastics now contaminate seafood, honey, milk, salt, water, soil, and air, even in remote regions such as the Arctic and Antarctic [7,9,10,10–14]. Recent studies have even detected microplastics in the human body, raising concerns about potential health risks [15].

The small size (< 5 mm) of these particles alone poses a risk to living organisms, as their dimensions are similar to cells and molecules. Micro and nanoplastic (size < 1 µm) particles (MNP) have been shown to accumulate in invertebrates and vertebrates (fish, birds, mammals) where they cause changes in feeding behaviour, growth, and development inhibition, hormonal disruption, gut dysbiosis and oxidative stress [7,14]. In fish, plastic particles absorbed through water, food or the trophic chain accumulate in the brain, ultimately modifying behavior [16,17]. Because plastic particles can interact with cell membranes and damage their integrity [18], it remains essential to investigate their effects on the nervous system. Oxidative stress and redox state are established biomarkers of acute toxicity and enable the identification of tissue-specific reactions to pollutants [19]. In addition, behavioral analyses represent sensitive and non-invasive indicators of toxicity and may reflect changes in fitness through disruption in copulatory, anti-predator or foraging behaviors [20,21].

Several studies have already synthesized and meta-analyzed the behavioral and toxic effects of microplastics on fish species [21–27]. Jacob calculated the percentage of affected endpoints as declared by the studies and indicated that most studies on behavioral, sensory, and neuromuscular functions or neurochemical outcomes showed some effect, mainly by smaller particles [21]. Overall negative or neutral effects have been reported for oxidative and acetylcholinesterase (AChE) biomarkers with influence of species, tissue, exposure routes, life stages, and microplastic types [22,27]. However, most previous meta-analyses aggregate multiple endpoints into a single overall model and examine moderators in separate models, without employing a multivariate, multilevel framework that can handle the correlated, hierarchical nature of the data.

Here, we report the results of a systematic review and meta-analysis of the effects of MNP on central nervous system outcomes of fish subjected to controlled experimental designs. We used a hierarchical random-effects model to account for data dependencies and to summarize experimental evidence on 13 outcomes related to neurochemistry and behavior of fish. We further investigated the potential influence of research authorship networks, fish species, and plastic type on the effect size estimates. In addition, we explored developmental stage, exposure duration, particle size, and concentration as potential sources of heterogeneity for each evaluated outcome. Given the substantial impact of exposure duration on the developmental stages, we also examined interactions between these moderators. Then, we assessed the overall validity of the results by analyzing publication bias (small study effects and time-lag bias) and identifying influential studies. Finally, we evaluated results discrepancies and similarities on a random effects multivariate analysis with a specified correlation matrix.

## 2. Material and Methods

The protocol for this review was pre-registered in Open Science Framework and is available at https://osf.io/ga9p4 [28]. This report adheres to the Preferred Reporting Items for Systematic Reviews and Meta-Analysis (PRISMA) 2020 guidelines [29,30]. Repository of data files: authors table, extraction guide, reporting quality assessment, particles composition, full list of outcomes, and codes are available at https://github.com/NNNESP/Nanofish_SR.

Relevant records were searched on June 27th, 2021, from PubMed, Web of Science, and Scopus. The search strategy (https://osf.io/px3vq) combined terms for population (fish, common fish names and fish species) and intervention (micro- or nanoplastic), with no further filters applied. Duplicate reports were identified and excluded using Rayyan [31], then randomly allocated to five groups by a developed Matlab tool (https://github.com/qkzan/PaperRandomizer).

### 2.1. Eligibility

Since plastic has a broad definition, we adopted the plastic definition described previously [32], which considers chemical composition, solid state (melting temperature Tm or glass transition temperature Tg > 20 °C), and solubility (<1 mg/L at 20 °C). Additionally, study selection was restricted to an environmental scope, as several polymers used in medical applications require further clarification regarding their chemical and thermal properties.

Research articles eligible were preclinical studies using adequate control groups (same model organism, same procedure) of fish exposed to virgin MNP under controlled conditions with behavioral or neurochemical outcomes. Records with no original data (reviews, letters, and comments) and non-interventional studies or studies that did not assess outcomes related to central nervous system function were excluded. Studies that investigated the central nervous system but only for gene expression, transcriptomic, proteomic analysis, ethoxyresorufin-O-deethylase to measure enzymatic activity, and Comet assays were excluded.

### 2.2. Selection

Report selection was done in two phases: pre-screening based on title/abstract and full text screening. In both phases, each paper was screened by two independent reviewers and discrepancies were resolved by a third reviewer. Rayyan was used to keep records and blind the decisions until each phase’s conclusion. Pre-screening reasons for exclusion were set for the type of study, population, or intervention. Additional exclusion reasons during full text screening were: absence of a control group or outcome of interest. Cohen’s Kappa was calculated to evaluate decision agreement on each independent group of researchers during study selection.

### 2.3. Extraction

The total number of studies were equally distributed between reviewers for data extraction. Qualitative extraction of studies included data for identification: first author name, DOI, url, authors, title, journal, year; particle characteristics: size, material, shape; animal information: species, sex, developmental stage and age at the start of MNP exposure; and experimental protocol: method of administration, dose/concentration, duration of exposure, time interval between last administration of particles and outcome assessment. Outcomes were selected for quantitative data extraction if at least 5 papers for the outcome were identified, except for speed, which was excluded due to its redundancy with distance data. Included neurochemical outcomes were acetylcholinesterase (AChE) activity, catalase (CAT) activity, superoxide dismutase (SOD) activity, glutathione peroxidase (GPx) activity, glutathione S-transferase (GST) activity, glutathione (GSH) content, total reactive oxygen species (ROS) levels (Hydrogen peroxide, ammonium molybdate, DCFDA) and lipid peroxidation (TBARS / MDA / LPO / CHP / FOX). Included behavioral outcomes were sensory-motor function evaluated by distance on light and dark tests, motor function evaluated by distance on free swimming tests, predatory performance, feeding time and speed during feeding.

In experiments where the age of the animals was not specified, evidence of reproduction was taken to indicate that they were adults. Prolarvae, larvae, fingerlings, and juveniles were grouped in the larvae and juveniles categories. If days post hatching or days post fertilization were provided, literature data were used for developmental stage classification (Table S1). Adult fish without sex specification were considered of unclear sex if the fish species was dioecious. The sexual system of each species was investigated using Fishbase, searching by species name and selecting the keyword “*reproduction”* on the “*Life cycle and mating behavior”* section [33]. When not explicitly stated, the frequency of administration was assumed to be once, the time interval between the last administration and testing less than 24 hours, and the water exposure under 24 hours as static administration. Partial or total water changes (10–100%) were considered semi-static.

Amino resins (aminoplasts) are condensation thermosetting polymers of formaldehyde with either urea or melamine [34]. Therefore, as melamine formaldehyde listed for one experiment [35] is an amino formaldehyde polymer (aminoplast) and five studies use aminoplasts of undisclosed composition [36–40], all of those were considered aminoplasts. Low-density polyethylene was grouped with polyethylene for quantitative analysis. It was not possible to access the size of the particle of one publication, in which the author confirmed by e-mail that the particles were smaller than 5 mm [35].

Quantitative data was extracted from text and tables or from figures using WebPlotDigitizer [41]. One paper presented individualized measurements for each animal, then the mean and standard deviation were calculated [42]. When repeated measurements of the same observational unit were presented, the longest exposure was selected [43]. Since light and dark tests had high variability in data presentation, data extraction priority was: first dark period, total dark period, and total distance [39,44–53].

If the number of independent observations was unclear, we calculated it from the degrees of freedom of anova results. Nonetheless, authors of 15 papers were contacted via email for further clarification on values extraction [35,45,46,51,54–62]. We received responses from 13 authors, while two studies were excluded from the meta-analysis due to data being presented as box plot and no response to the contact [56,57].

### 2.4. Quality report assessment

The analysis of overall reporting quality was adapted from the reporting standards for rigorous study design proposed previously [63]. The presence of descriptions for the following items was evaluated: (1) random allocation of animals to treatment groups; (2) a priori sample size estimation method; (3) post-exposure data exclusion or animal inclusion criteria; (4) blinding for allocation of animals or outcome assessor during experiment. If at least one experiment within a study explicitly addressed each of these methodological aspects, the study was assigned a corresponding ‘yes’ for that criterion. Initially, 10% of studies were accessed by all the reviewers for fine-tuning of evaluation criteria. Each of the other papers included was screened by two independent reviewers, and discrepancies were resolved in group discussion. Plots were created with robvis [64].

### 2.5. Modularity analysis

To investigate authorship dependence as a possible influence on studies, the network of researchers was determined [65]. Briefly, a co-authorship network map was created with VOSviewer with full counting, not ignoring documents with a large number of authors, and no threshold for the number of publications or citations per author [66]. The output was saved as a GML file and uploaded to Gephi 0.10. with random decomposition, weights from edges, resolution = 1, and software’s default settings [67]. Then, the study cluster was manually assigned based on the cluster of its authors.

### 2.6. Effect measures and synthesis methods

Each outcome of interest was analyzed individually. Effect sizes were determined with standardized mean differences (SMD) by escalc function [68]. Based on data structure, with multiple effect sizes by study and shared controls, a correlated and hierarchical random effects model with robust variance estimation, with a restricted maximum likelihood estimation method was used for overall results estimation and heterogeneity evaluation [69,70]. Heterogeneity was evaluated by Cochran’s Q, total I², and σ^2^ for each random effect level [71]. The hierarchical/nested model accounts for the inter-dependencies between data, where data are grouped inside a higher level of another group. In this meta-analysis the individual effect sizes are grouped inside a control group identifier (ES cluster), which is a part of the study. Additional random effects terms considered were species, type of plastic, and study cluster. Likelihood ratio tests with model estimates by the maximum likelihood method were used to evaluate the relevance of including these terms for each database.

Same control groups correlations in effect sizes were controlled with ES cluster random term inclusion. Correlated sampling errors due to repeated use of the same animals were controlled with a variance-covariance matrix calculation with correlation coefficients of 0.5 [71]. The ESid was included considering protocol variations of each independent effect size [71]. Variance components were inspected with profile likelihood plots to verify if the models were overfitted. Based on data structure, the outcomes feeding time and speed during feeding had a simplified random effects formula without ES cluster.

Cook’s distance was used to examine what effect the deletion of one study has on the fitted values of all studies simultaneously as influential cases analysis [70]. Cases characteristics were used to identify patterns for inference of potential moderators [70]. Also, sensitivity analysis of the overall effect was conducted with variation of correlation coefficient (⍴ = 0, 0.25, 0.75, and 0.9) and leave-one-out analysis on the study level. Publication bias for small study-effects, when small studies (with small sample sizes) tend to report large effect sizes, was evaluated by the PET-PEESE method together with the inclusion of year (centered) as a moderator for time-lag bias - decline in reported effect sizes over time [71]. Sensitivity analysis of small study-effects was performed with ‘effective sample size’ that accounts for unbalanced sampling [72]. GPx activity, GST activity, Feeding time and Speed during feeding did not converge for the combined moderator analysis for publication bias, so we analysed it separately for those outcomes.

Meta-regression was performed for outcomes with at least 10 data clusters and studies. A correlated hierarchical mixed-effects model with robust estimation was used to partially explain the heterogeneity of data. Based on the question, data availability, and relevance of each parameter, the chosen moderators were developmental stage, exposure duration, particle size, and concentration. Developmental stage was added as a categorical variable, and the numeric ones were standardized as follows: exposure duration in days, mean particle size in micrometers, and concentration in mg/L. When concentration data were only provided as the number of particles/m³ or similar scales in the study, it was considered unclear and the study was dropped from meta-regression. All analyses were conducted in R 4.4.1 [73]. Finally, with multivariate analysis it is possible to represent correlations through a correlated multivariate model with robust variance estimation. In order to do that, *vcalc* correlation matrix was established with the study as cluster, outcome as type, and control and treated groups specifications. Results were compared with the individual outcomes analysis with and without publication bias correction.

### 2.7. Deviations from protocol

The following outcomes were excluded during extraction: morphometric measures, survival, speed. Also, meta-analysis methods were changed from random effects to correlated hierarchical random effects to account for data dependencies. Meta-regression would be used for outcomes with at least 10 experiments. However, we evaluated at least 10 studies since true independent observations were on study level for the hierarchical model.

## 3. Results

### 3.1. Study selection

In total, 4.389 records were collected from the databases. After duplicates removal, 2633 records went through title and abstract screening, 1717 were excluded by type of study, 346 by population, and 189 by intervention reasons, and so 379 were sought for retrieval (Fig. 1). One study was not retrieved [74], and 378 went through full text screening. Substantial agreement of 81.85% ± 1.74% (CI95%: 78.4%-85.3%) was achieved between reviewers (Cohen’s Kappa).

**Figure 1.**
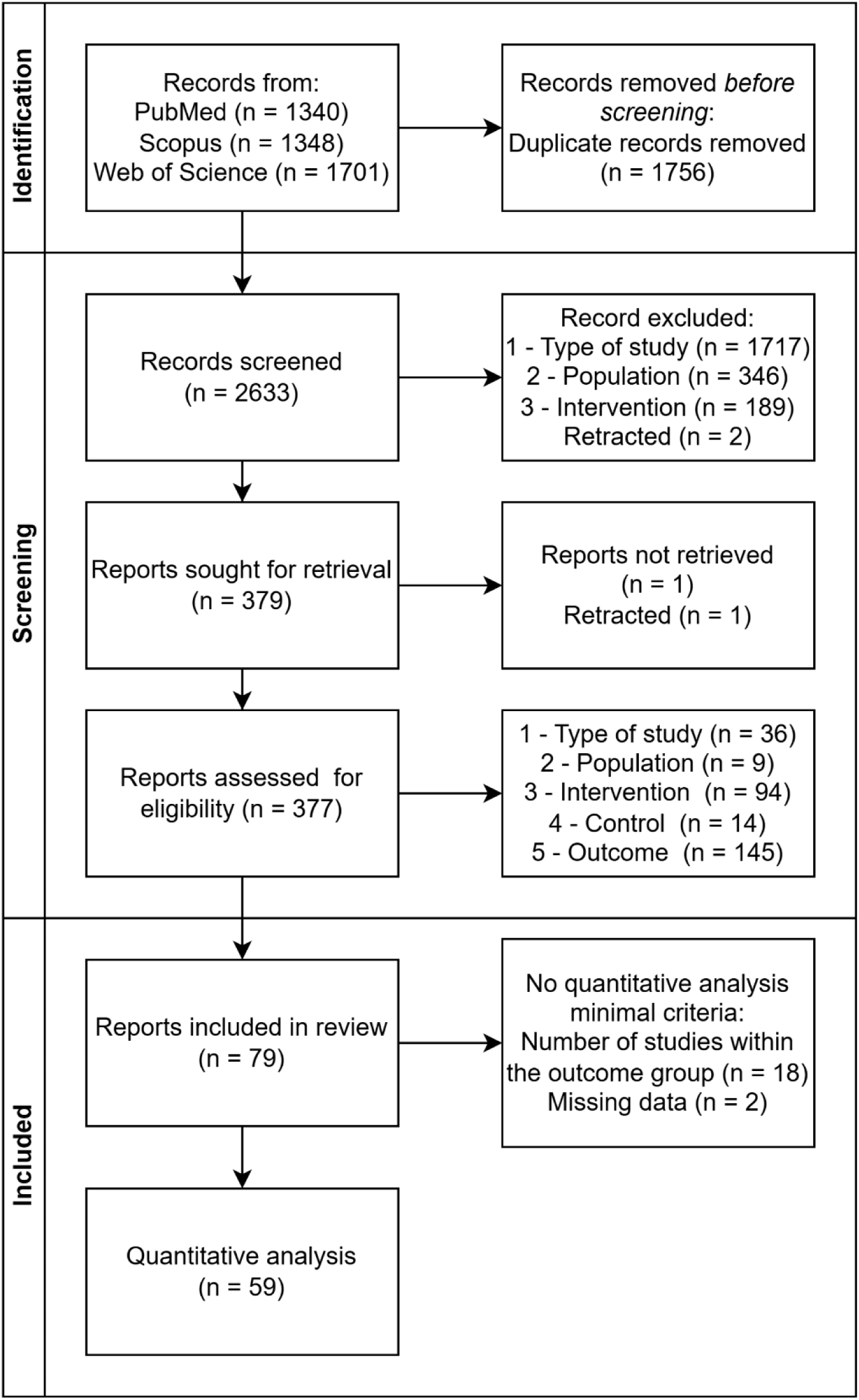
Flow chart diagram of the studies collection and selection procedure.

During full text screening, two retracted papers were identified with PubPeer and excluded by type of study [75,76]. Among the remaining studies, 80 presented outcomes related to behavior or brain neurochemistry and were included in the qualitative synthesis (Table 1). Reasons for exclusion at the full text screening phase were because criteria were not met for type of study (n = 36), population of interest (n = 9), type of intervention (n = 94), proper control (n = 14), outcome of interest (n = 145, Fig. 1). Cohen’s Kappa between researchers for this phase was 63.15% ± 3.98% (CI95%: 55.34%-70.97%).

**Table 1:**
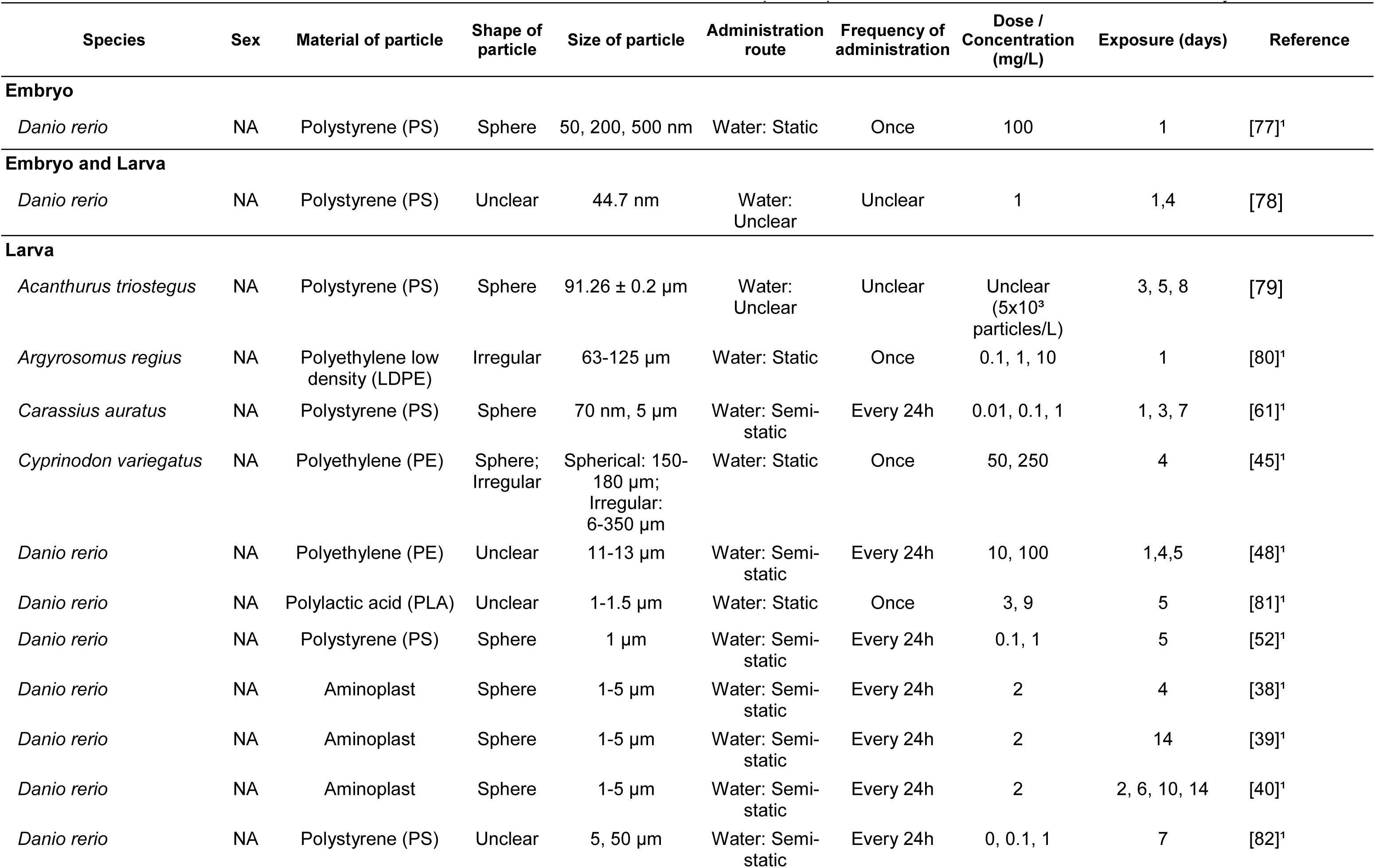
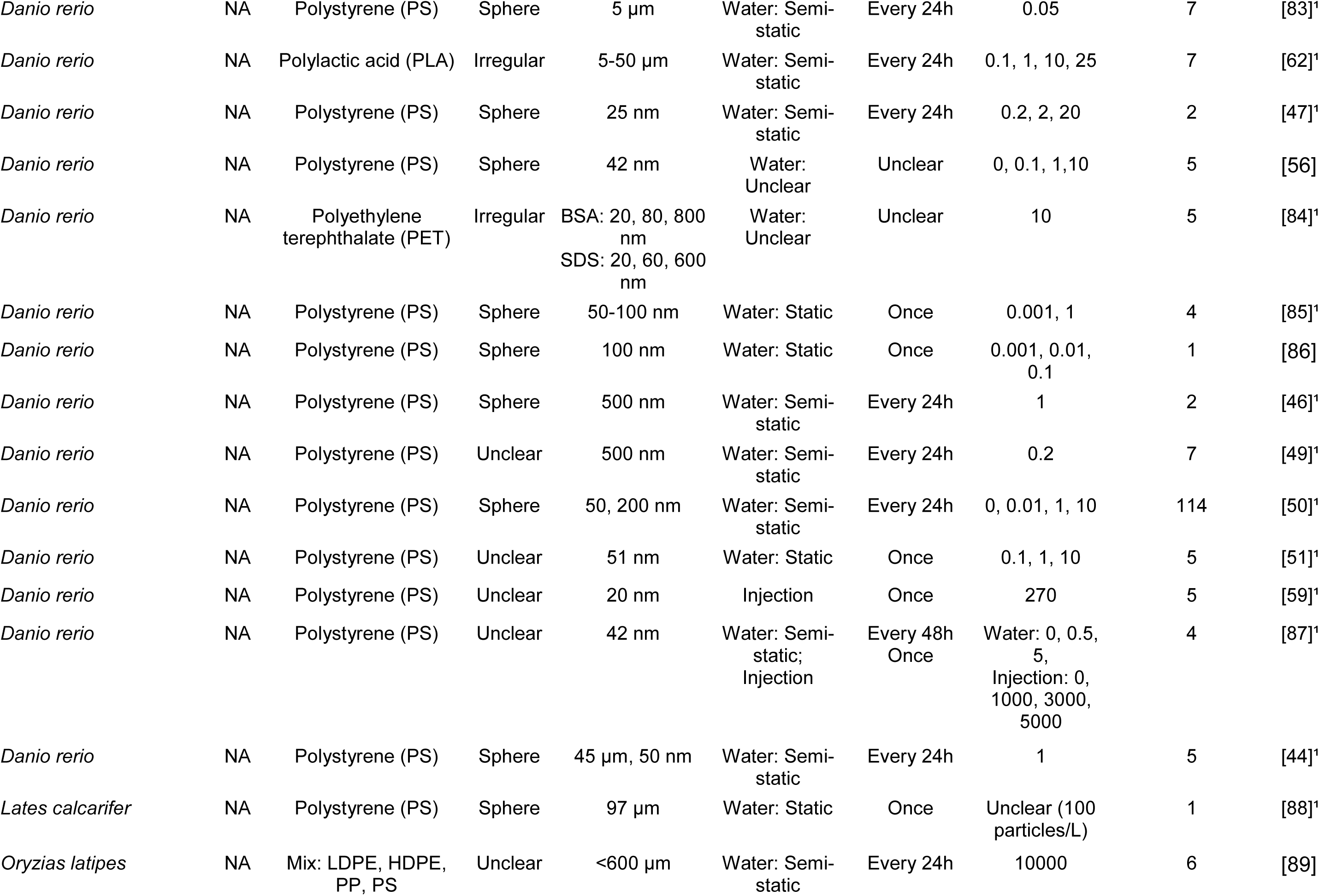
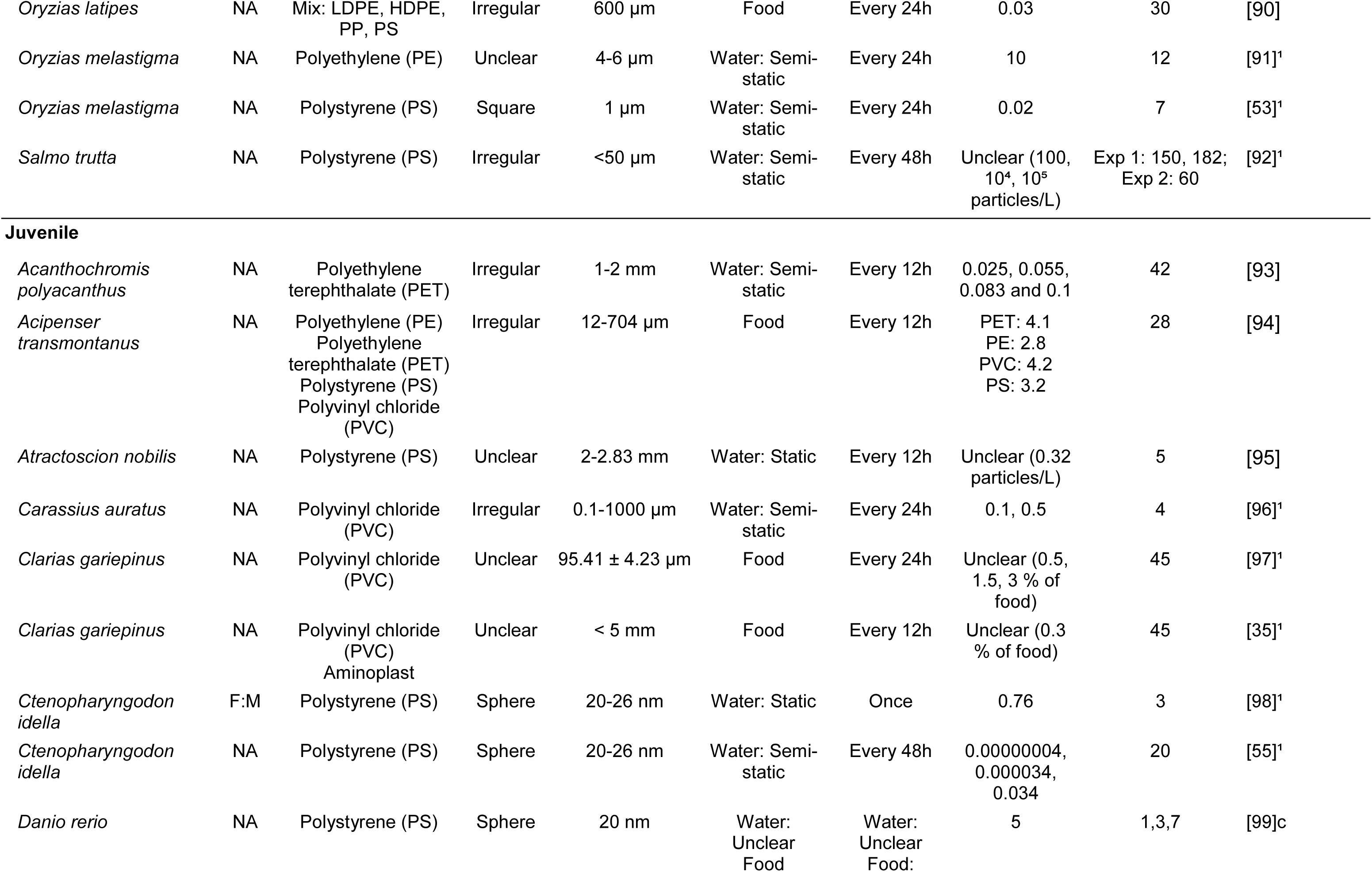
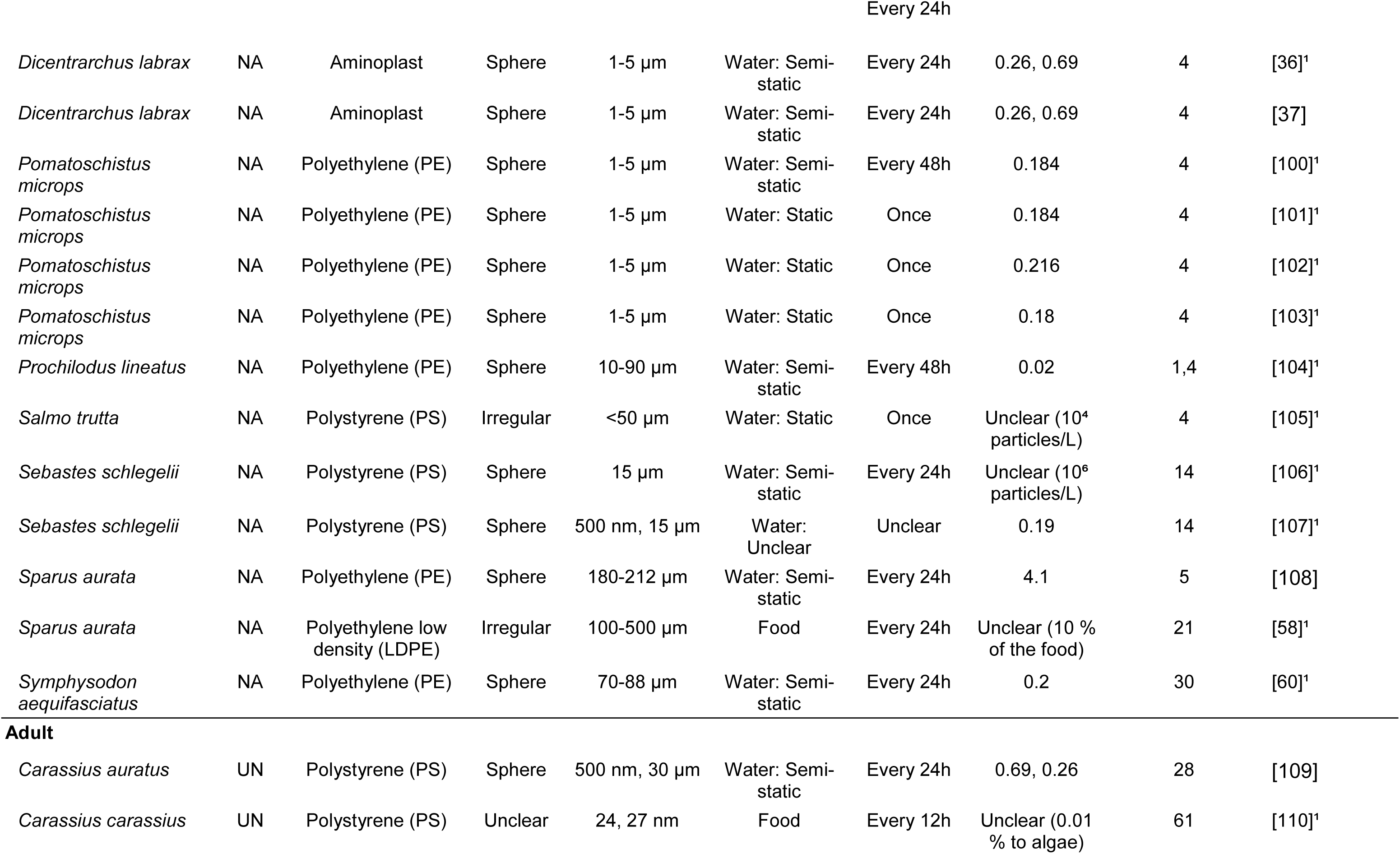
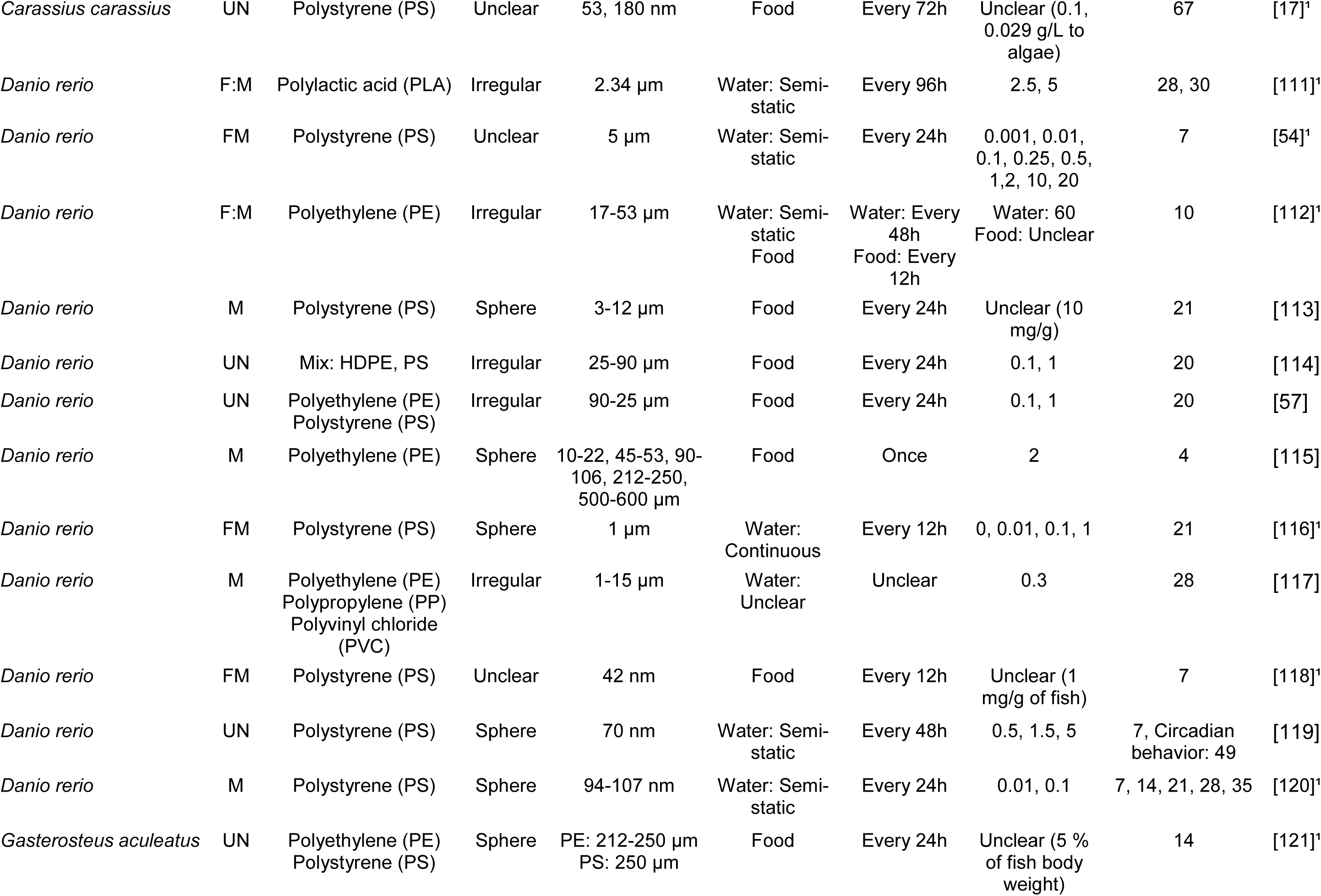
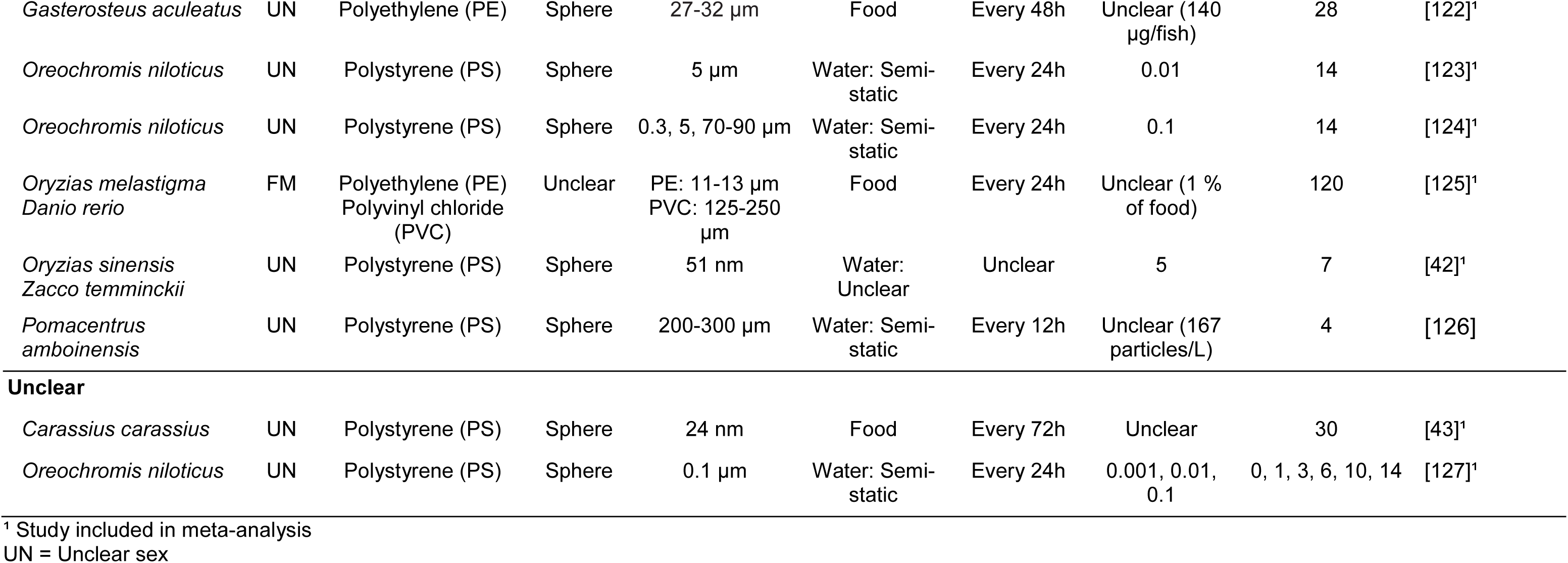
Studies included in this review that evaluated effects of micro or nano sized plastic particles on behavior or brain neurochemistry of fish.

Decisions from both phases together had 79.76% ± 2.39%, CI95% from 75.07% to 84.45% of agreement between researchers, indicating substantial agreement. Of the 80 reports initially included, one was retracted during the course of this study and was therefore later excluded from the analysis. The minimal criteria for quantitative meta-analysis was met for 59 reports (Fig. 1).

### 3.2. Study characteristics

Publication year of the reports ranged from 2012 to 2021, with exponential growth, considering that data collection was in June 2021 (Fig. 2a, Table 1). Nine journals published at least 3 of the included reports, with most of them from Journal of Hazardous Materials (13 reports) and Aquatic Toxicology (10 reports, Fig. 2b.). Cluster analysis from the 397 authors and 1434 edges identified 46 researcher groups with minimal crossings between them (modularity index = 0.960). Authors with higher links were Jérôme Cachot (23 weight links, 5 reports, cluster 1), Christelle Clérandeau (21 weight links, 4 reports, cluster 1), Florane Le Bihanic (19 weight links, 3 reports, cluster 1) and Guilherme Malafaia (19 weight links, 5 reports, cluster 2, Fig. 2d and <u>VOSviewer Online</u>). All four of them had a mean year of publication between 2019.8 and 2020.8 (Fig. 2d). The author with more reports published was Lúcia Guilhermino, with 6 reports (12 weight links, cluster 6, Fig. 2d).

**Figure 2.**
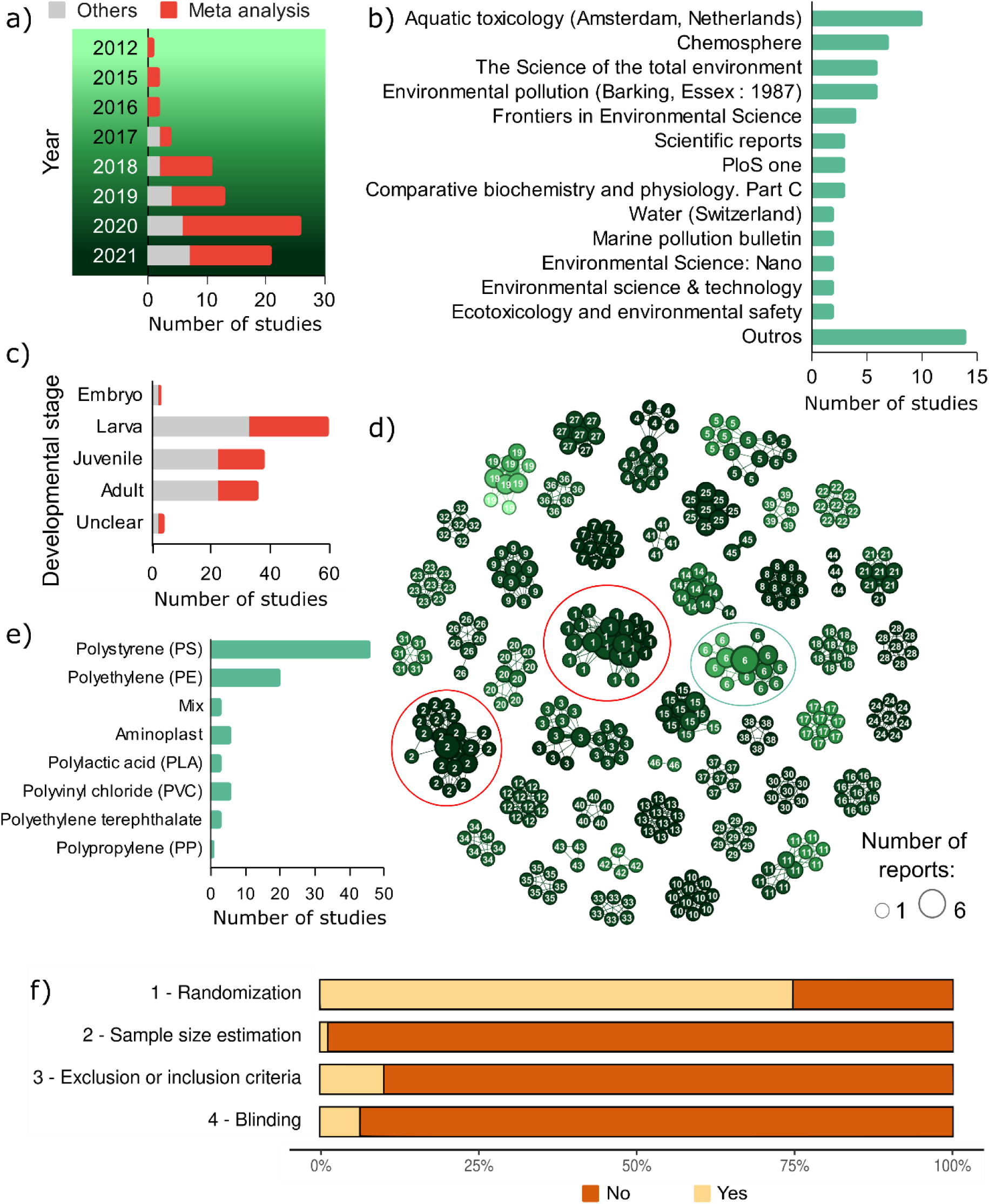
Scientometry and quality of report of studies that investigate micro or nano plastic particles effects on brain and behavior. a) Histogram of publications by year of all included reports (gray) or the ones that achieved criteria to meta analysis (red). b) Number of studies published in each journal. c) Number of studies with fish on each developmental stage during outcome assessment. d) Co-authorship network analysis of researchers. Two recent research groups had higher links and reports (red circle). The research group with the author with more reports: Lúcia Guilhermino with 6 reports (green circle). Size of nodes indicate number of reports by author and colour indicates the average publication year of the studies published by each researcher, according to the background color in a). Line thickness represents correlations between researchers. e) Number studies that included each plastic type. f) Quality of report summary with percentage of the studies presenting at least once each feature.

Fish species were native from all over the world, and they had a diverse habitat, mostly from unknown population trends and low concern for extinction (Table 2). Nevertheless, *Acipenser transmontanus* is a vulnerable species, and *Prochilodus lineatus*, *Sebastes schlegelii* and *Zacco temminckii* are not evaluated as stated at the IUCN Red List of Threatened Species [128]. Those species appeared in only one or two studies. The only fish species with more than 4 studies was *Danio rerio,* which accounted for 45.6% (36/79) of the reports.

**Table 2.**
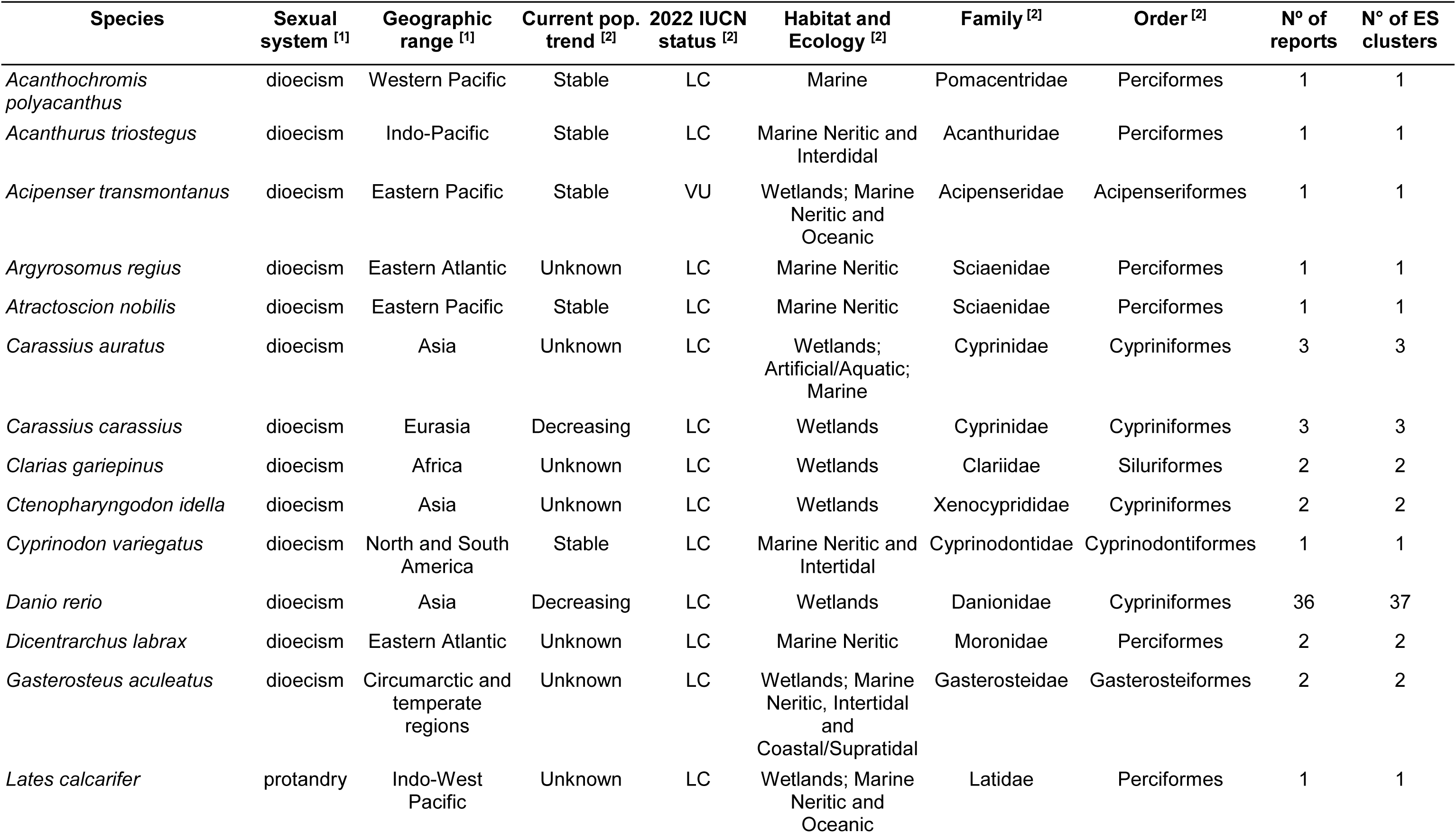
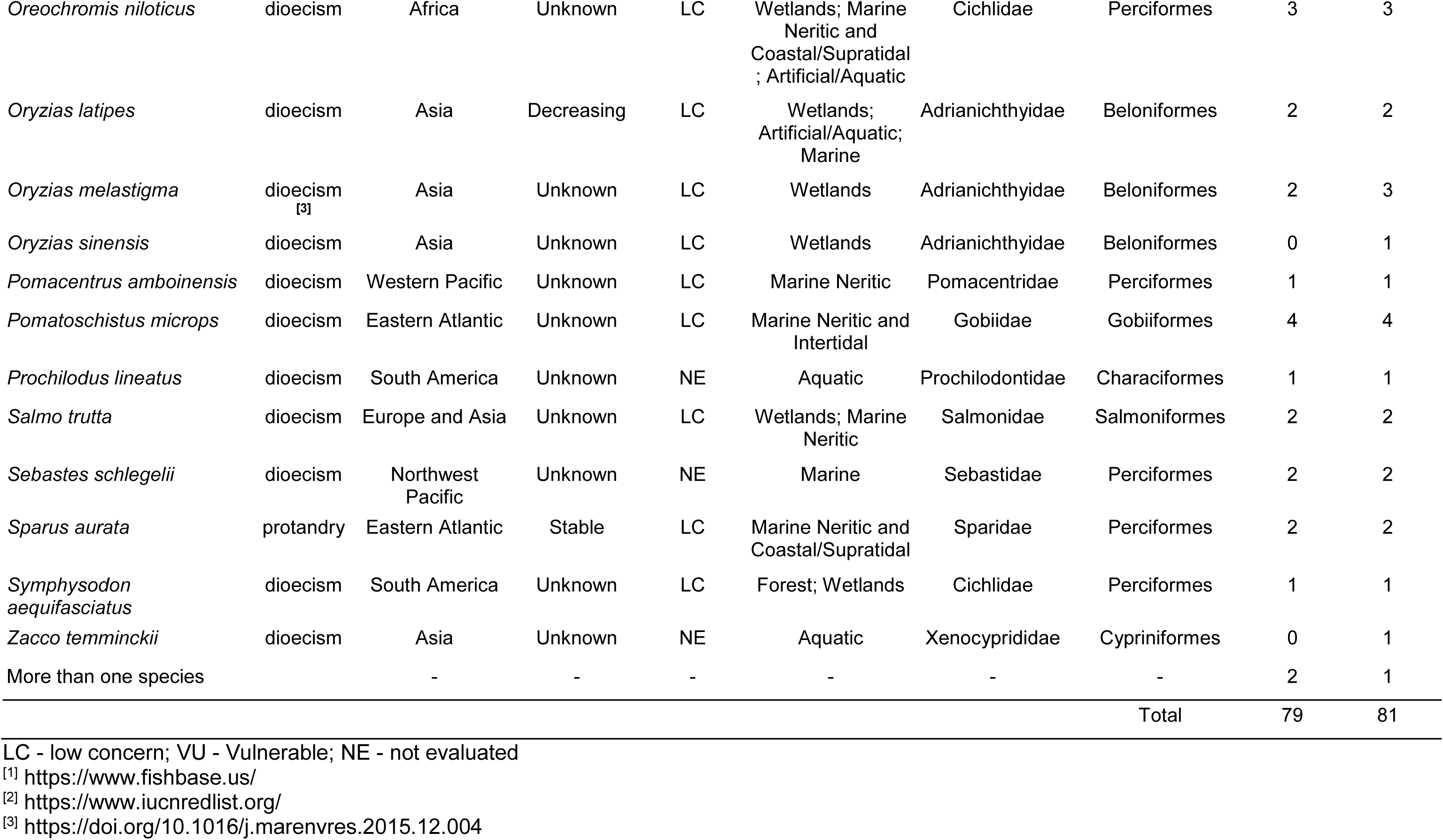
Species used in the reports and their characteristics with occurrence on the reports and ES clusters.LC - low concern; VU - Vulnerable; NE - not evaluated.

| Species | Sexual system <sup>[1]</sup> | Geographic range <sup>[1]</sup> | Current pop. trend <sup>[2]</sup> | 2022 IUCN status <sup>[2]</sup> | Habitat and Ecology <sup>[2]</sup> | Family <sup>[2]</sup> | Order <sup>[2]</sup> | Nº of reports | Nº of ES clusters |
| --- | --- | --- | --- | --- | --- | --- | --- | --- | --- |
| *Acanthochromis polyacanthus* | dioecism | Western Pacific | Stable | LC | Marine | Pomacentridae | Perciformes | 1 | 1 |
| *Acanthurus triostegus* | dioecism | Indo-Pacific | Stable | LC | Marine Neritic and Intertidal | Acanthuridae | Perciformes | 1 | 1 |
| *Acipenser transmontanus* | dioecism | Eastern Pacific | Stable | VU | Wetlands; Marine Neritic and Oceanic | Acipenseridae | Acipenseriformes | 1 | 1 |
| *Argyrosomus regius* | dioecism | Eastern Atlantic | Unknown | LC | Marine Neritic | Sciaenidae | Perciformes | 1 | 1 |
| *Atractoscion nobilis* | dioecism | Eastern Pacific | Stable | LC | Marine Neritic | Sciaenidae | Perciformes | 1 | 1 |
| *Carassius auratus* | dioecism | Asia | Unknown | LC | Wetlands; Artificial/Aquatic; Marine | Cyprinidae | Cypriniformes | 3 | 3 |
| *Carassius carassius* | dioecism | Eurasia | Decreasing | LC | Wetlands | Cyprinidae | Cypriniformes | 3 | 3 |
| *Clarias gariepinus* | dioecism | Africa | Unknown | LC | Wetlands | Clariidae | Siluriformes | 2 | 2 |
| *Ctenopharyngodon idella* | dioecism | Asia | Unknown | LC | Wetlands | Xenocyprididae | Cypriniformes | 2 | 2 |
| *Cyprinodon variegatus* | dioecism | North and South America | Stable | LC | Marine Neritic and Intertidal | Cyprinodontidae | Cyprinodontiformes | 1 | 1 |
| *Danio rerio* | dioecism | Asia | Decreasing | LC | Wetlands | Danionidae | Cypriniformes | 36 | 37 |
| *Dicentrarchus labrax* | dioecism | Eastern Atlantic | Unknown | LC | Marine Neritic | Moronidae | Perciformes | 2 | 2 |
| *Gasterosteus aculeatus* | dioecism | Circumarctic and temperate regions | Unknown | LC | Wetlands; Marine Neritic, Intertidal and Coastal/Supratidal | Gasterosteidae | Gasterosteiformes | 2 | 2 |
| *Lates calcarifer* | protandry | Indo-West Pacific | Unknown | LC | Wetlands; Marine Neritic and Oceanic | Latidae | Perciformes | 1 | 1 |
| *Oreochromis niloticus* | dioecism | Africa | Unknown | LC | Wetlands; Marine<br>Neritic and<br>Coastal/Supratidal<br>; Artificial/Aquatic | Cichlidae | Perciformes | 3 | 3 |
| *Oryzias latipes* | dioecism | Asia | Decreasing | LC | Wetlands;<br>Artificial/Aquatic;<br>Marine | Adrianichthyidae | Beloniformes | 2 | 2 |
| *Oryzias melastigma* | dioecism<br>[3] | Asia | Unknown | LC | Wetlands | Adrianichthyidae | Beloniformes | 2 | 3 |
| *Oryzias sinensis* | dioecism | Asia | Unknown | LC | Wetlands | Adrianichthyidae | Beloniformes | 0 | 1 |
| *Pomacentrus amboinensis* | dioecism | Western Pacific | Unknown | LC | Marine Neritic | Pomacentridae | Perciformes | 1 | 1 |
| *Pomatoschistus microps* | dioecism | Eastern Atlantic | Unknown | LC | Marine Neritic and<br>Intertidal | Gobiidae | Gobiiformes | 4 | 4 |
| *Prochilodus lineatus* | dioecism | South America | Unknown | NE | Aquatic | Prochilodontidae | Characiformes | 1 | 1 |
| *Salmo trutta* | dioecism | Europe and Asia | Unknown | LC | Wetlands; Marine<br>Neritic | Salmonidae | Salmoniformes | 2 | 2 |
| *Sebastes schlegelii* | dioecism | Northwest<br>Pacific | Unknown | NE | Marine | Sebastidae | Perciformes | 2 | 2 |
| *Sparus aurata* | protandry | Eastern Atlantic | Stable | LC | Marine Neritic and<br>Coastal/Supratidal | Sparidae | Perciformes | 2 | 2 |
| *Symphysodon*<br>*aequifasciatus* | dioecism | South America | Unknown | LC | Forest; Wetlands | Cichlidae | Perciformes | 1 | 1 |
| *Zacco temminckii* | dioecism | Asia | Unknown | NE | Aquatic | Xenocyprididae | Cypriniformes | 0 | 1 |
| More than one species |  | - | - | - | - | - | - | 2 | 1 |
| Total |  |  |  |  |  |  |  | 79 | 81 |

[1] https://www.fishbase.us/
[2] https://www.iucnredlist.org/
[3] https://doi.org/10.1016/j.marenvres.2015.12.004

Of the species reported, only two – *Lates calcarifer* and *Sparus aurata –* are protandric, and in the three relevant reports they were used as larvae or juveniles. All remaining species are dioecious, so the sex should be reported for adult individuals (Tables 2 and S1). From 24 reports with adult or unclear developmental stage of dioecious species, 14 were unclear for sex, 4 were only male, 4 reported separate experiments for females and males, and two had mixed sexes. Additionally, one study reported mixed proportional sex distribution of *Ctenopharyngodon idella,* although they were used as juveniles [98]. Most reports evaluated outcomes on larvae (31/79 included, 26/59 meta-analysed) followed by juveniles (22/79, 16/59) and adult animals (22/79, 14/59); one investigated both embryos and larvae and one only embryos; 2 studies did not report any information on the developmental stage of the animals (Fig. 2c, Table 1).

Plastic composition of the particles investigated in the included reports were polystyrene (PS), polyethylene (PE), amino formaldehyde polymer (Aminoplast), polylactic acid (PLA), polyvinyl chloride (PVC), polyethylene terephthalate (PET) and polypropylene (PP) (Fig. 2e). PS particles were the most common plastic composition studied (49/79, 34/59), followed by PE (25/79, 16/59). Nine reports studied particles with more than one plastic composition, and 3 of them with mixtures (Table 3). The shape of the particles was irregular, sphere or square, with a concentration range from 0.04 ng/L to 10g/L and size range from 20 nm to 1000 µm.

**Table 3.**
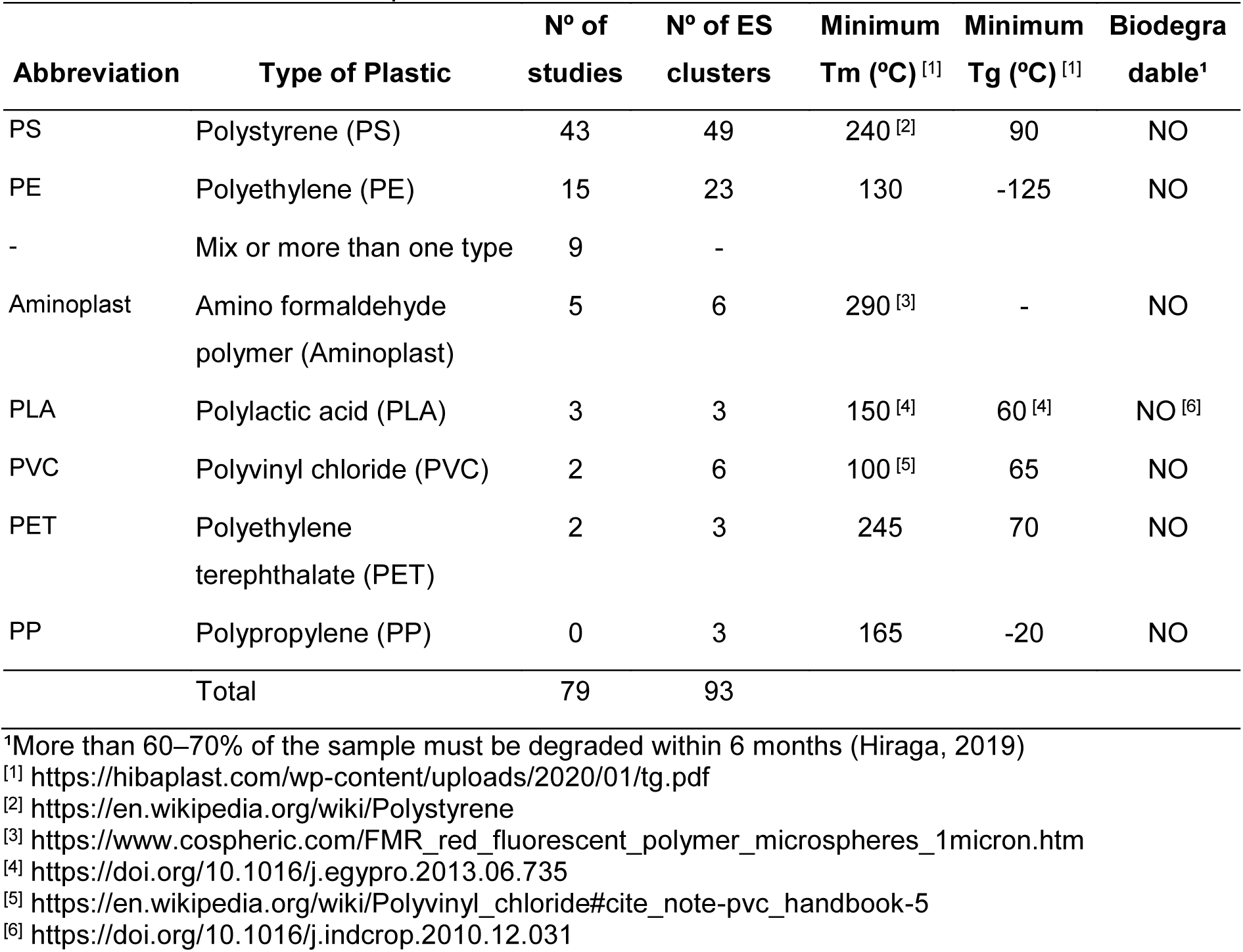
Synthetic polymer or semi-synthetic polymer properties and frequency of occurrence in the included reports.

| Abbreviation | Type of Plastic | N° of studies | N° of ES clusters | Minimum Tm (°C) <sup>[1]</sup> | Minimum Tg (°C) <sup>[1]</sup> | Biodegradable<sup>1</sup> |
| --- | --- | --- | --- | --- | --- | --- |
| PS | Polystyrene (PS) | 43 | 49 | 240 <sup>[2]</sup> | 90 | NO |
| PE | Polyethylene (PE) | 15 | 23 | 130 | -125 | NO |
| - | Mix or more than one type | 9 | - |  |  |  |
| Aminoplast | Amino formaldehyde polymer (Aminoplast) | 5 | 6 | 290 <sup>[3]</sup> | - | NO |
| PLA | Polylactic acid (PLA) | 3 | 3 | 150 <sup>[4]</sup> | 60 <sup>[4]</sup> | NO <sup>[6]</sup> |
| PVC | Polyvinyl chloride (PVC) | 2 | 6 | 100 <sup>[5]</sup> | 65 | NO |
| PET | Polyethylene terephthalate (PET) | 2 | 3 | 245 | 70 | NO |
| PP | Polypropylene (PP) | 0 | 3 | 165 | -20 | NO |
| Total |  | 79 | 93 |  |  |  |

<sup>1</sup>More than 60–70% of the sample must be degraded within 6 months (Hiraga, 2019)
<sup>[1]</sup> https://hibaplast.com/wp-content/uploads/2020/01/tg.pdf
<sup>[2]</sup> https://en.wikipedia.org/wiki/Polystyrene
<sup>[3]</sup> https://www.cospheric.com/FMR_red_fluorescent_polymer_microspheres_1micron.htm
<sup>[4]</sup> https://doi.org/10.1016/j.egypro.2013.06.735
<sup>[5]</sup> https://en.wikipedia.org/wiki/Polyvinyl_chloride#cite_note-pvc_handbook-5
<sup>[6]</sup> https://doi.org/10.1016/j.indcrop.2010.12.031

### 3.3. Quality of report in studies

Overall quality of report in studies was poor, with only one study [108] scoring ‘yes’ for 3 of the 4 criteria evaluated, while 10 studies scored ‘yes’ for 2 of them, and 19 did not report any of the evaluated criteria (Fig. S1). Randomization was informed at least once for 59 studies, exclusion or inclusion criteria for 8, blinding for 5 and sample size estimation for only one study (Fig. 2f, Fig. S1). Overall reviewers’ agreement was high at 84.83% (Cohen’s Kappa: randomization = 81.77%, sample size estimation = 100%, exclusion or inclusion criteria = 53.47% and blinding = 88.15%). Considering the general poor reporting quality of the included studies, it was not reasonable to apply the SYRCLE’s risk of bias assessment tool [129].

### 3.4. Quantitative synthesis

Meta-analyses were performed with the 59 studies that met the criteria of data availability and presence of an outcome reported in at least 5 studies (Table 1). The protocols employed in the studies show high variability, covering all developmental stages, including one study in embryos (Table 1, Fig. 3a). In general, exposure duration ranged from 1 to 182 days, with most reports within 1 to 15 days (Fig. 3b). Larval and juvenile experiments were mostly under 7 days; only 5/26 reports for larvae and 7/16 for juveniles had longer exposure to the treatment (Fig. 1a, Table 1). The minimal exposure duration for adult animals was 7 days, and half of the reports covered experiments with exposures longer than 15 days (Fig. 3a, Table 1).

**Figure 3.**
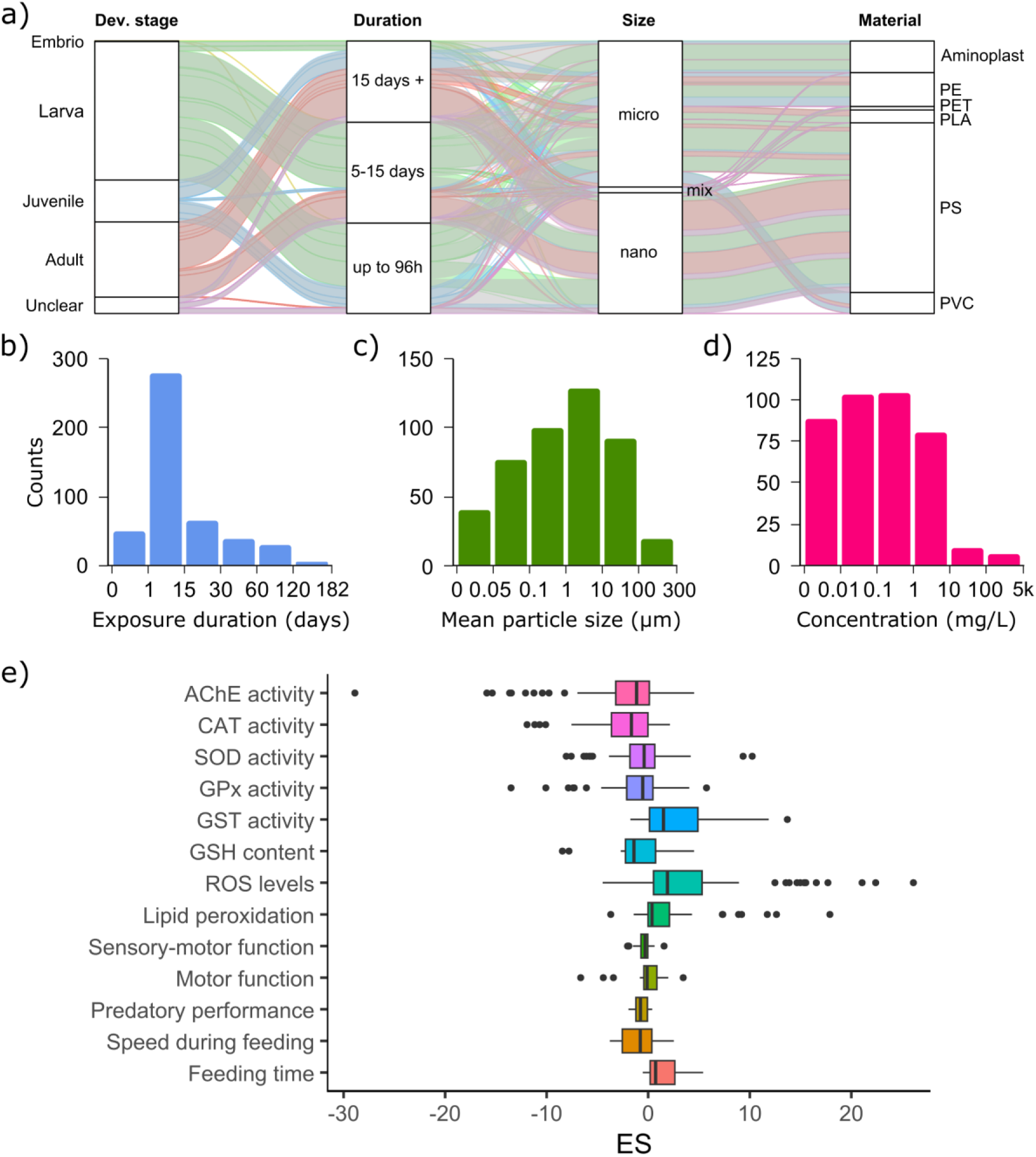
Descriptive plots of the effect sizes (ES) included in the meta-analysis. a) Alluvial plot showing protocol variability, color coded by developmental stage. b-d) Histograms of exposure duration (b), mean particle size (c) and concentration (d) of the included ES. e) Boxplot of the ES values by outcome.

Micro and nano size categories were well represented on reports of all three developmental stages; mean particle size ranged from 20 nm to 300 µm, with one study with mixed sized particles ([96], Fig. 3c). Plastic types for larva reports were Aminoplast, PE, PET, PLA and PS; juvenile: Aminoplats, PE, PS and PVC; and adult: PE, PLA, PS and PVC (Fig. 3a). Concentration ranged from 3 x 10^-8^ to 5000 mg/L, with the extreme values reported in the two studies in *Danio rerio* larvae with administration via injection ([59,87], 2020, Fig. 3d).

Considering the extracted data, adding random effects for study cluster, species, or plastic type did not significantly improve model fit for any of the outcomes examined and were not included for further analysis.

#### 3.4.1. Neurochemical outcomes

##### 3.4.1.1. AChE activity

The meta-analysis comprised 83 comparisons from 50 data clusters of 25 independent studies. There was no difference between treatment and control groups (ES = -0.67 ± 0.38, CI95% = -1.45 to 0.11, prediction interval -4.42 to 3.08), with high heterogeneity (I²_total_ = 83.96%; Q_82_ = 310.23, p < 0.001) and variance explained by each component of σ²_Study_ = 2.94, σ²_EScluster_ = 0.07, σ²_ESid_ = 0.14 (Fig. 4). The Cook’s distance highlighted the Umamaheswari et al. (2021) [120] and Chagas et al (2021) [111] as influential reports. Observing features extracted from the papers, there was no other obvious moderator to be included other than the ones already selected for meta-regression. Umamaheswari et al. (2021) [120] has very high ES values, even over 50 for one of the groups and Santos et al. (2021) [40] very high variance over 40 for one of the groups. Even so, sensitivity analysis of the overall estimate indicates robust results with no difference between control and exposure groups except for Estrela et al (2021) [98] and Chagas et al (2021) [111] removal. Publication bias results (F_2_,_3.8_ = 19.12, p = 0.010) indicate the absence of time-lag bias (t_6.26_ = 1.19, p = 0.278), but the presence of small-study effect (t_2.29_ = -6.94, p = 0.014, Fig. S2). This means smaller studies have larger effect sizes. Sensitivity analysis for small-study effect using effective sample size indicates absence of small-study effect (F_2,8.59_ = 1.01, p = 0.403; t_12_ = -1.50, p = 0.159). Nevertheless, even considering the possibility of small-study effect influence, the conditional overall estimate sustained the result with no difference between microplastic exposure and control groups (ES = -0.18 ± 0.34, CI95% = -0.89 to 0.53, prediction interval -3.28 to 2.92), with high heterogeneity (QE_80_ = 214.50 p < 0.001). The meta-regression comprised 67 comparisons from 38 data clusters of 20 independent studies (QE_57_ = 248.20, p < 0.001). No moderator included was significant.

**Figure 4.**
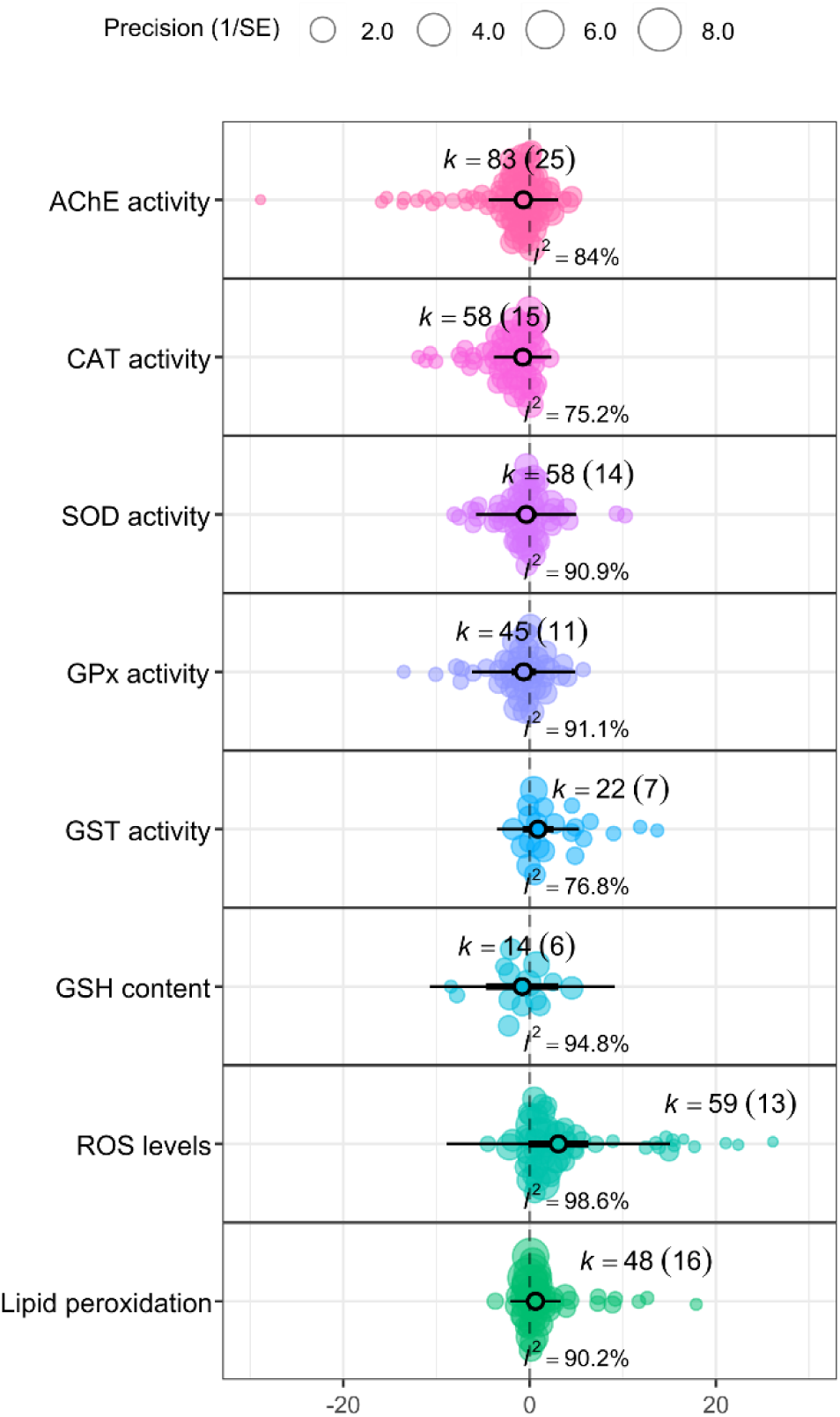
Orchard plots of the overall effects for biochemical outcomes. Filled circle = individual effect sizes (ES) scaled by sample size, open circle = point estimate with associated 95% confidence interval (thick black horizontal line) and prediction intervals (thin black horizontal line), k = number of ES (number of reports).

##### 3.4.1.2. CAT activity

The meta-analysis comprised 58 comparisons from 25 data clusters of 15 independent studies. There was no difference between treatment and control groups (ES = -0.74 ± 0.37, CI95% = -1.54 to 0.06, prediction interval -3.83 to 2.35), with high heterogeneity (I²_total_ = 75.24%; Q_57_ = 182.62, p < 0.001) and variance explained by each component of σ²_Study_ = 1.45, σ²_EScluster_ = 0.00, σ²_ESid_ = 0.48 (Fig. 4). The Cook’s distance highlighted the Umamaheswari et al. (2021) as an influential report [120]. Observing features extracted from the papers, there was no other obvious moderator to be included other than the ones already selected for meta-regression. Even so, sensitivity analysis of the overall estimate indicates robust results with no difference between control and exposure groups except for Rios-Fuster et. al. (2021) [58] removal with rho 0.25 and 0.5. Publication bias results (F_2_,_2.79_ = 15.58, p = 0.030) indicate the absence of time-lag bias, but the presence of small-study effect (t_1.92_ = -6.29, p = 0.027). Sensitivity analysis for small-study effect with effective sample size indicates absence of small-study effect (F_2,4.59_ = 0.607, p = 0.583; t_7.12_ = 0.17, p = 0.866). Nevertheless, even considering the possibility of small-study effect influence, the conditional overall estimate sustained the result with no difference between groups (ES = 2.42 ± 0.46, CI95% = 1.17 to 3.67, prediction interval -0.10 to 4.94), with high heterogeneity (QE_80_ = 78.60, p = 0.020).

The meta-regression comprised 48 comparisons from 19 data clusters of 10 independent studies. Longer exposure of fish larvae to plastic particles resulted in lower catalase (CAT) activity compared to shorter exposures (larvae: 0.39 ± 0.07, t_2.52_ = 5.78, p = 0.016, CI95% = 0.15 to 0.64). No other moderator included was significant. Residual heterogeneity was still high (QE_42_ = 92.22, p < 0.001) with similar variance explained between studies and ESid (σ²_Study_ = 0.51, σ²_EScluster_ = 0.00, σ²_ESid_ = 0.53).

##### 3.4.1.3. SOD activity

The meta-analysis comprised 58 comparisons from 23 data clusters of 14 independent studies. There was no difference between treatment and control groups (ES = -0.35 ± 0.51, CI95% = -1.49 to 0.77, prediction interval -5.75 to 5.04), with high heterogeneity (I²_total_ = 90.92%; Q_57_ = 398.52, p < 0.001) and variance explained by each component of σ²_Study_ = 1.63, σ²_EScluster_ = 0.00, σ²_ESid_ = 4.15 (Fig. 4).The Cook’s distance highlighted the Umamaheswari et al. (2021) as an influential report [120]. Observing features extracted from the papers, there was no other obvious moderator to be included other than the ones already selected for meta-regression. Sensitivity analysis of the overall estimate indicates robust results with no difference between control and exposure groups. Publication bias results (F_2,2.79_ = 0.14, p = 0.871) indicate the absence of time-lag bias and small-study effect. The meta-regression comprised 40 comparisons from 14 data clusters of 9 independent studies. Only one study with adult animals was dropped from the model. No moderator included was significant.

##### 3.4.1.4. GPx activity

The meta-analysis comprised 45 comparisons from 21 data clusters of 11 independent studies. There was no difference between treatment and control groups (ES = -0.64 ± 0.58, CI95% = -1.98 to 0.69, prediction interval -6.19 to 4.90), with high heterogeneity (I²_total_ = 91.12%; Q_44_ = 302.35, p < 0.001) and variance explained by each component of σ²_Study_ = 1.41, σ²_EScluster_ = 0.00, σ²_ESid_ = 4.08 (Fig. 4).The Cook’s distance highlighted the Umamaheswari et al. (2021) as an influential report [120]. Observing features extracted from the papers, there was no other obvious moderator to be included other than the ones already selected for meta-regression. Even so, sensitivity analysis of the overall estimate indicates robust results with no difference between control and exposure groups. Publication bias results indicate the absence of time-lag bias (F_1_,_2.2_ = 0.10, p = 0.774) and small-study effect (F_1_,_2.17_ = 1.79, p = 0.304).

##### 3.4.1.5. GST activity

The meta-analysis comprised 22 comparisons from 14 data clusters of 7 independent studies. There was no difference between treatment and control groups (ES = 0.90 ± 0.67, CI95% = -0.74 to 2.55, prediction interval -3.51 to 5.32), with high heterogeneity (I²_total_ = 76.80%; Q_21_ = 71.66, p < 0.001) and variance explained by each component of σ²_Study_ = 2.00, σ²_EScluster_ = 0.00, σ²_ESid_ = 0.78 (Fig. 4). The Cook’s distance highlighted the Umamaheswari et al. (2021) as an influential report [120]. Observing features extracted from the papers, there was no other obvious moderator to be included other than the ones already selected for meta-regression. Even so, sensitivity analysis of the overall estimate indicates robust results with no difference between control and exposure groups. Although it was not possible to evaluate time-lag bias due to convergence issues, publication bias results indicate the presence of small-study effect (F_1_,_1.1_ = 585.19, p = 0.019, t_1.1_ = 24.19, p = 0.02). Sensitivity analysis for small-study effect with effective sample size indicates absence of small-study effect (F_2,4.59_ = 0.069, p = 0.814; t_2.23_ = -0.26, p = 0.815). Nevertheless, even considering the possibility of small-study effect influence, the conditional overall estimate sustained the result with no difference between groups (ES = 0.07 ± 0.39, CI95% = -0.88 to 1.03, prediction interval -2.30 to 2.44), with moderate heterogeneity (QE_20_ = 30.69, p = 0.059).

##### 3.4.1.6. GSH content

The meta-analysis comprised 14 comparisons from 9 data clusters of 6 independent studies. There was no difference between treatment and control groups (ES = -0.80 ± 1.51, CI95% = -4.69 to 3.10, prediction interval -10.76 to 9.16), with high heterogeneity (I²_total_ = 94.80%; Q_13_ = 76.35, p < 0.001) and variance explained by each component of σ²_Study_ = 12.14, σ²_EScluster_ = 0.36, σ²_ESid_ = 0.09 (Fig. 4). There were no influential studies. Sensitivity analysis of the overall estimate indicates robust results with no difference between control and exposure groups. Publication bias results (F_2_,_0.79_ = 1.82, p = 0.507) indicate the absence of time-lag bias and small-study effect.

##### 3.4.1.7. Total ROS levels

The meta-analysis comprised 59 comparisons from 24 data clusters of 13 independent studies. There was no difference between treatment and control groups (ES = 3.08 ± 1.47, CI95% = -0.11 to 6.28, prediction interval -8.92 to 15.08), with high heterogeneity (I²_total_ = 98.6%; Q_58_ = 605.99, p < 0.001) and variance explained by each component of σ²_Study_ = 26.43, σ²_EScluster_ = 0.00, σ²_ESid_ = 1.73 (Fig. 4). The Cook’s distance highlighted the Umamaheswari et al. (2021) [120] and Sökmen et al. (2020) [59] as influential reports. Observing features extracted from the papers, there was no other obvious moderator to be included other than the ones already selected for meta-regression. Even so, sensitivity analysis of the overall estimate indicates robust results with no difference between control and exposure groups except for Estrela et al. (2021) [98] removal with rho ⩽ 0.75 or full database with an absence of correlation (rho = 0). Publication bias results (F_2_,_3.05_ = 8.75, p = 0.054) indicate the absence of time-lag bias, but the presence of small-study effect (t_2.4_ = 3.87, p = 0.045). Sensitivity analysis for small-study effect with effective sample size indicates absence of small-study effect (F_2,4.25_ = 0.521, p = 0.623; t_4.07_ = 0.17, p = 0.350). Nevertheless, even considering the possibility of small-study effect influence, the conditional overall estimate sustained the result with no difference between groups (ES = -1.16 ± 1.32, CI95% = -4.09 to 1.78, predicton interval -10.09 to 7.78), with high heterogeneity (QE_56_ = 422.56, p = 0.001). The meta-regression comprised 56 comparisons from 22 data clusters of 11 independent studies. No moderator included was significant. The Wald test could not be computed because the variance-covariance matrix of the contrast was not positive definite, also we did not compute a reduced database model since there was no exclusive study for any developmental stage.

##### 3.4.1.8. Lipid peroxidation

The meta-analysis comprised 48 comparisons from 25 data clusters of 16 independent studies. There was no difference between treatment and control groups (ES = 0.63 ± 0.33, CI95% = -0.08 to 1.34, prediction interval -2.06 to 3.32), with high heterogeneity (I²_total_ = 90.20%; Q_47_ = 175.62, p < 0.001) and variance explained by each component of σ²_Study_ = 1.44, σ²_EScluster_ = 0.00, σ²_ESid_ = 0.04 (Fig. 4). The Cook’s distance highlighted the Umamaheswari et al. (2021) [120] and Chagas et. al. (2021) [111] as influential reports. Observing features extracted from the papers, there was no other obvious moderator to be included other than the ones already selected for meta-regression. Sensitivity analysis of the overall estimate indicates robust results with no difference between control and exposure groups except for Santos et. al. (2020) [38] removal for any correlation value. Publication bias results (F_2,2.46_ = 2.94, p = 0.222) indicate the absence of time-lag bias and small-study effect. The meta-regression comprised 31 comparisons from 17 data clusters of 10 independent studies. No moderator included was significant.

#### 3.4.2. Behavior

##### 3.4.2.2. Sensory-motor function

The meta-analysis comprised 30 comparisons from 16 data clusters of 14 independent studies, all in larvae. Treatment animals travel shorter distances than control groups in sensory-motor test (ES = -0.42 ± 0.18, CI95% = -0.82 to -0.03, prediction interval -1.94 to 1.09), with high heterogeneity (I²_total_ = 90.79%; Q_29_ = 294.31, p < 0.001) and variance explained by each component of σ²_Study_ = 0.26, σ²_EScluster_ = 0.03, σ²_ESid_ = 0.16 (Fig. 5). The Cook’s distance highlighted the Zhang et al. (2020) as an influential report [87]. Observing features extracted from the papers, there was no other obvious moderator to be included other than the ones already selected for meta-regression. Sensitivity analysis of the overall estimate indicates controversial results, with the exclusion of 5 reports reinforcing the result and the 9 others without statistical significance of the treatment effect. Publication bias results (F_2_,_3.81_ = 0.80, p = 0.511) indicate the absence of time-lag bias and small-study effect. The meta-regression comprised 30 comparisons from 16 data clusters of 14 independent studies. There were only studies on the larval stage, so the developmental stage was not included as a moderator. No included moderator was significant.

**Figure 5.**
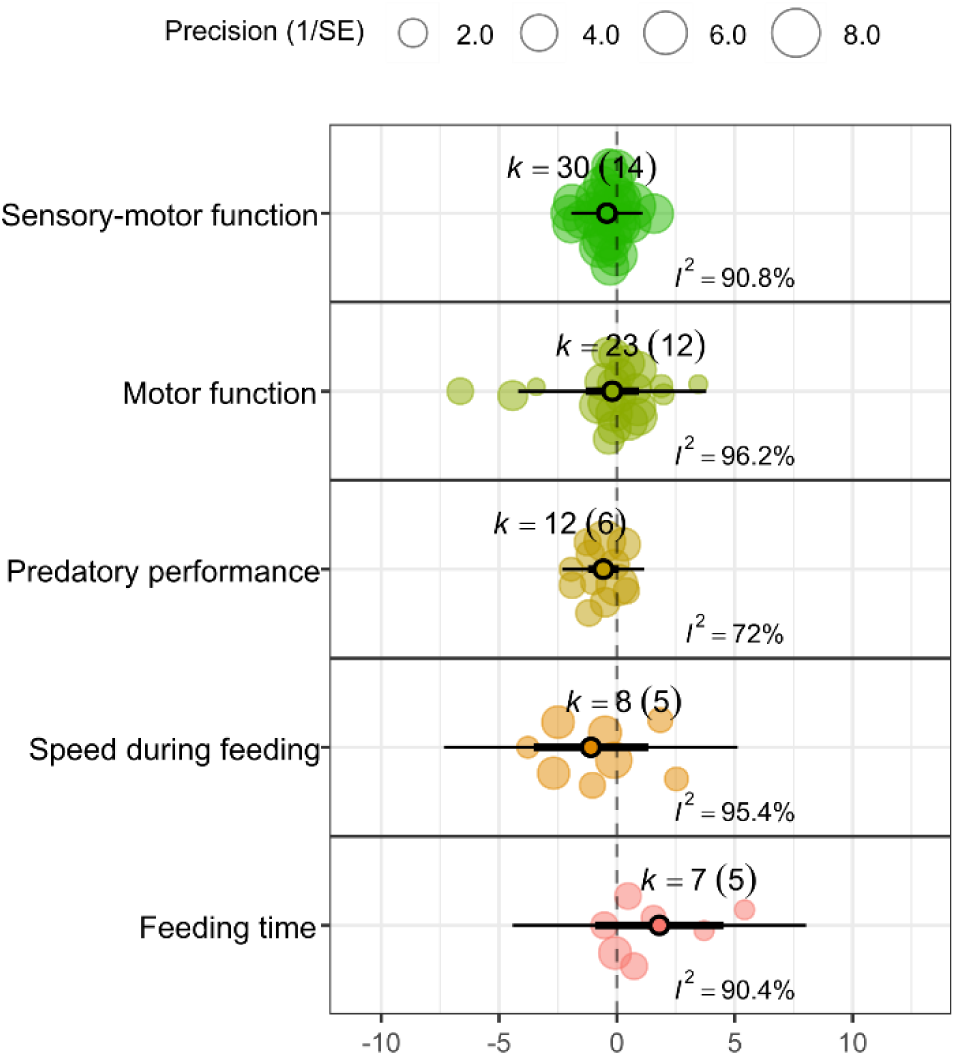
Orchard plots of the overall effects for behavioral outcomes. Fsh larvae exposed to micro or nano plastic travel shorter distances as compared to controls on sensory-motor function tasks. Filled circle = individual effect sizes (ES) scaled by sample size, open circle = point estimate with associated 95% confidence interval (thick black horizontal line) and prediction intervals (thin black horizontal line), k = number of ES (number of reports).

##### 3.4.2.1. Motor function

The meta-analysis comprised 23 comparisons from 17 data clusters of 12 independent studies. There was no difference between treatment and control groups (ES = -0.20 ± 0.51, CI95% = -1.32 to 0.92, prediction interval -4.19 to 3.79), with high heterogeneity (I²_total_ = 96.22%; Q_22_ = 211.87, p < 0.001) and variance explained by each component of σ²_Study_ = 2.77, σ²_EScluster_ = 0.00, σ²_ESid_ = 0.25 (Fig. 5). The Cook’s distance highlighted the de Oliveira et al. (2021) as an influential report [81]. Observing features extracted from the papers, there was no other obvious moderator to be included other than the ones already selected for meta-regression. Sensitivity analysis of the overall estimate indicates robust results with no difference between control and exposure groups. Publication bias results (F_2,1.58_ = 0.43, p = 0.707) indicate the absence of time-lag bias and small-study effect. The meta-regression comprised 17 comparisons from 12 data clusters of 10 independent studies. No included moderator was significant.

##### 3.4.2.3. Predatory performance

The meta-analysis comprised 12 comparisons from 10 data clusters of 6 independent studies, only in larva or juvenile fish. There was no difference between treatment and control groups (ES = -0.57 ± 0.25, CI95% = -1.23 to 0.08, prediction interval -2.32 to 1.16), with high heterogeneity (I²_total_ = 72.00%; Q_11_ = 31.33, p < 0.001) and variance explained by each component of σ²_Study_ = 0.09, σ²_EScluster_ = 0.00, σ²_ESid_ = 0.28 (Fig. 5). Sensitivity analysis of the overall estimate indicates robust results with no difference between control and exposure groups. Publication bias results (F_2,1.38_ = 1.04, p = 0.529) indicate the absence of time-lag bias and small-study effect.

##### 3.4.2.4. Speed during feeding

The meta-analysis comprised 8 comparisons from 5 data clusters of 5 independent studies, the random formula was ∼1 | Study/ESid. There was no difference between treatment and control groups (ES = -1.10 ± 0.84, CI95% = -3.53 to 1.32, prediction interval -7.33 to 5.12), with high heterogeneity (I²_total_ = 95.40%; Q_7_ = 121.92, p < 0.001) and variance explained by each component of σ²_Study_ = 1.34 and σ²_ESid_ = 2.65 (Fig. 5). Sensitivity analysis of the overall estimate indicates no difference between control and exposure groups for all leave-one-out and correlation coefficient variation analysis. Publication bias results indicate an absence of time-lag bias (F_1,1.44_ = 0.16, p = 0.739) and small-study effect (F_1,2.04_ = 0.0001, p = 0.994).

##### 3.4.2.5. Feeding time

The meta-analysis comprised 7 comparisons from 5 data clusters of 5 independent studies, the random formula was ∼1 | Study/ESid. There was no difference between treatment and control groups (ES = 1.80 ± 0.97, CI95% = -0.93 to 4.53, prediction interval -4.43 to 8.03), with high heterogeneity (I²_total_ = 90.37%; Q_6_ = 26.46, p < 0.001) and variance explained by each component of σ²_Study_ = 3.67, σ²_ESid_ = 0.28 (Fig. 5). Sensitivity analysis of the overall estimate indicates robust results with no difference between control and exposure groups. Publication bias results indicate an absence of time-lag bias (F_1,1.56_ = 9.34, p = 0.125) and presence of small-study effect (F_1_,_2.13_ = 22.44, p = 0.037, t_2.13_ = 4.74, p = 0.037). Sensitivity analysis for small-study effect with effective sample size corroborates with presence of small-study effect (F_1,1.74_ = 25.08, p = 0.050; t_1.74_ = 5.01, p = 0.050). Considering the small-study effect influence, the conditional overall estimate sustained the result with no difference between groups (ES = -1.85 ± 0.71, CI95% = -5.13 to 1.42, prediction interval -5.64 to 1.94), without heterogeneity (QE_5_ = 7.02, p = 0.219).

#### 3.4.3. Multivariate analysis

The multivariate analysis with the 59 studies and 13 outcomes, 467 observations, indicates an increased lipid peroxidation and reduced sensory-motor function in MNP exposed fish (lipid peroxidation: ES = 0.75 ± 0.34, CI95% = 0.02 to 1.47; sensory-motor function: ES = -0.49 ± 0.17, CI95% = -0.85 to -0.12, Fig. 6a). However, this result was not consistent with publication bias correction inclusion (lipid peroxidation: t_5.32_ = 0.82, p = 0.448; sensory-motor function: t_6.5_ = -0.45, p = 0.666, Fig. 6b). Publication bias analysis results (F_13,12.97_ = 0.43, p = 0.707) indicate absence of time-lag bias for all outcomes and presence of small-study effects for AChE activity, GST activity and predatory performance (AChE: t_2.3_ = -6.55, p = 0.016; GST: t_1.13_ = 14.85, p = 0.031; predatory performance: t_1.68_ = -6.52, p = 0.035). As effective sample size accounts for unbalanced sampling, publication bias sensitivity was evaluated, and no outcome detected neurotoxic effects or publication bias. Moderator analysis did not converge with robust variance estimation.

**Figure 6.**
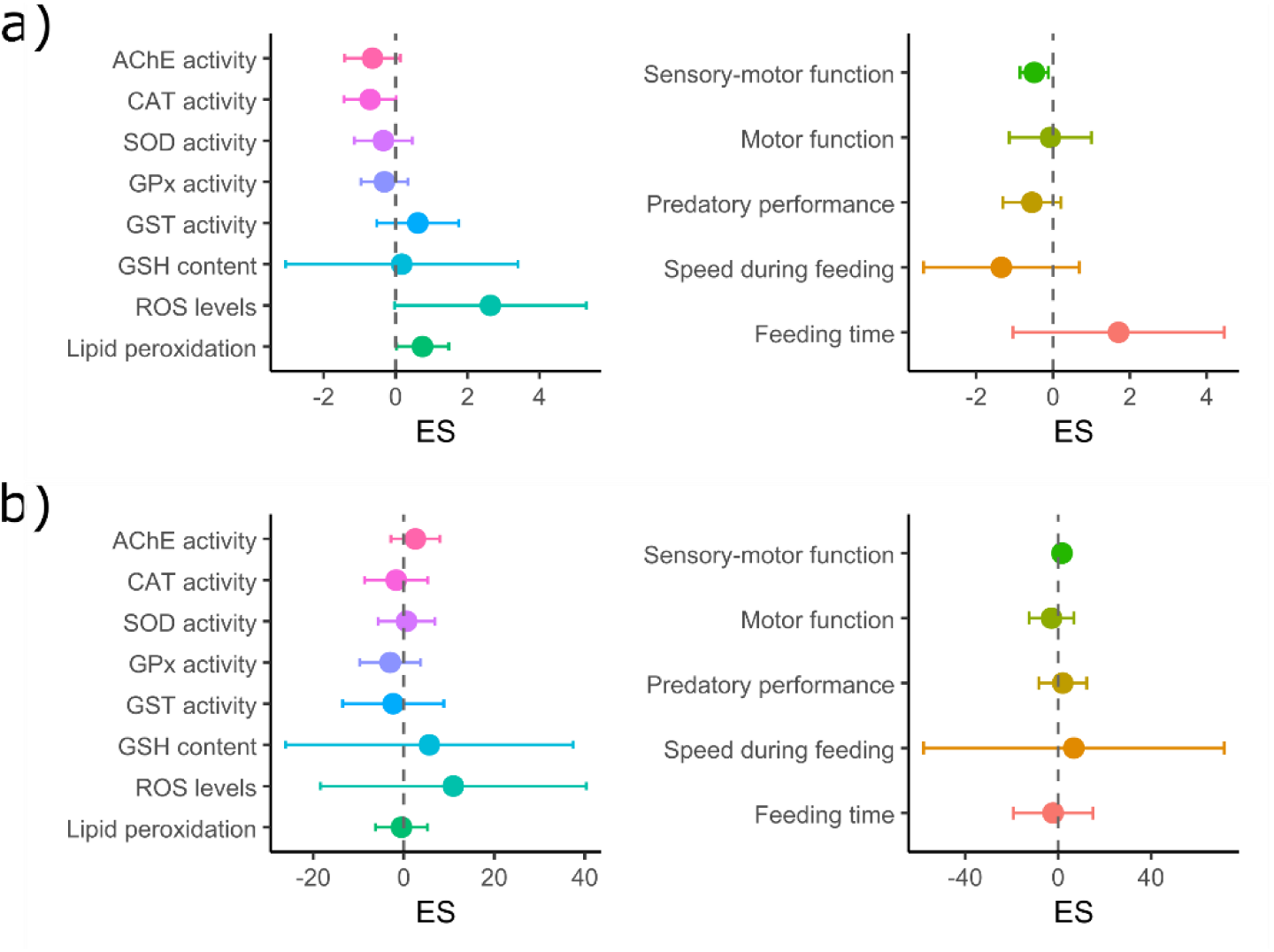
Multivariate analysis effect sizes (ES) for the 13 outcomes. (a) Increased lipid peroxidation and reduced sensory-motor function in micro or nano plastic exposed fish. (b) Publication bias corrected estimates. Filled circle = point estimate with associated 95% confidence interval.

## 4. Discussion

This systematic review and meta-analysis summarized evidence from 59 studies that investigated the effects of virgin MNP on 13 neurological endpoints in fish, with outcomes ranging from neurochemical biomarkers to behavioral performance. When data dependencies were properly handled with a hierarchical random effects model, no consistent effects of MNP exposure were observed for all but one of the outcomes. Sensory-motor function was the only outcome with a statistically significant summary effect; all included studies were conducted exclusively in larvae, and showed that MNP exposure shortened the traveled distance when compared to controls. However, this small effect (-0.42 [CI95% -0.82; -0.03) was deemed fragile considering the sensitivity analyses, and the prediction interval ranged from -1.94 to 1.09, meaning the effect in future studies could be negligible or even in the opposite direction.

Multivariate analysis with robust variance estimation could be used to estimate the effect size of neurochemical and behavioral and, yielding results that are mostly consistent with those obtained from the hierarchical random effects analysis. Although for multivariate analysis lipid peroxidation was significant, the confidence interval was 0.02 to 1.47, while for the hierarchical random effects it was -0.08 to 1.34, which is a very close range. The sensory-motor function showed a similar reduced travel distance in the exposed group. However, the multivariate modelling confirmed the overall absence of consistent effects induced by exposure to MNP once small-study effects were taken into account.

These results challenge the narrative that MNP are toxic to fish and affect several biological functions, as reported in previous reviews [21–27,130]. Our more conservative approach of accounting for shared controls and nested comparisons within studies suggests that many of the previously claimed effects in meta-analytical syntheses may reflect statistical artefacts instead of true neurotoxicity. On the other hand, the substantial heterogeneity observed across studies (I^2^ > 75%, mostly above 90% for most outcomes) could not be satisfactorily explained by the variables we tested as moderators, indicating that other, as-yet-unidentified factors could account for the between-study variance.

A rigorous assessment of the quality of the primary studies included in a meta-analysis is of utmost importance for properly evaluating and interpreting its results. An analysis of several systematic reviews of animal experiments showed that studies lacking randomization, blinded outcome assessment, and allocation concealment reported markedly inflated effect sizes [131]. Unfortunately, reporting of key methodological details was poor in the included studies, which hindered us from evaluating the underlying risk of bias, as most aspects would be rated as unclear for most of the studies. Only one study reported three of the four criteria we assessed (randomization, blinding, exclusion criteria, and sample size estimation), while nearly one-third failed to report any of them. This lack of transparency regarding methodological details and bias-reducing measures is another reason to interpret the results of this review with caution, as it is impossible to determine whether the authors did not report bias-reducing measures or whether they did not implement them. We cannot exclude the possibility that, if it was possible to assess study quality, different results could be obtained by restricting the meta-analyses to the highest-quality studies. Therefore, we strongly encourage future investigators to fully adhere to reporting checklists, such as the ARRIVE 2.0 guideline for in vivo studies [132], since more transparent reporting will bolster our confidence in the resulting evidence.

Potential biases were identified, particularly small-study effects, although time-lag bias was not detected for most outcomes. Small-study effects inflated estimates for AChE, CAT and GST activities, total ROS level, and feeding time. Although selective reporting likely amplified individual findings, this did not result in a consistent neurotoxic signal at the field level, as data dependencies were accounted for in the meta-analytical model.

This review has several limitations. First, database searches were conducted in 2021, and the field has grown rapidly since then, so updated searches may yield additional relevant studies. Second, changes to the pre-registered protocol were required during the review process (e.g., exclusion of some outcomes, shift in meta-analytic approach) due to unforeseen data structures, which introduced some degree of analytic flexibility. Third, the lack of a clear experimental unit identification in many studies complicated variance estimation and could have affected effect size precision. Importantly, only direct effects of MNPs were considered; transgenerational and ecological effects (e.g., through the food web or predator-prey interactions) were not evaluated and may underestimate the overall impact on fish fitness and survival. Furthermore, restricting the studies to virgin polymer particles enhances internal consistency but limits the ecological realism.

Despite these limitations, the study has important strengths. It covers a broad range of species, mitigating ecological bias and improving generalizability. It also focused on exposure via feeding routes, which closely mimic real-world scenarios, and compiled data in a reproducible format available for further community reanalysis and updates. The use of a hierarchical meta-analytic model allowed better handling of nested data and shared controls, often present in ecotoxicology studies.

To enhance future research in this area, we suggest large scale use of ARRIVE guidelines for reporting animal research and propose inclusion of at least the following minimal information in publications: (a) exact MNP dosage in mg/L or enough data for conversion, (b) physicochemical parameters of the exposure and test tanks, (c) use of dosages relevant for environmentally realistic exposures and (d) clear statement of the independent experimental unit and dependencies on observational units. These details are essential to enable reproducibility, comparison, and risk assessment across species and exposure contexts.

Future directions should focus on addressing key knowledge gaps, such as the neurotoxic effects of weathered or environmentally aged microplastics, which may differ significantly from virgin particles in surface chemistry and biological reactivity. Also, polystyrene particles were the most common (43/80) plastic composition studied, although in the environment PE [133,134], PP [135], and poliamide [136] are the most abundant types reported and should be investigated more frequently. Another example of a key knowledge gap was that a large proportion of studies did not differentiate between sexes, which hindered the identification of sex-specific effects of particles. Furthermore, greater attention is needed to evaluate reproductive outcomes and predator-prey interactions, behaviors that can also influence fitness in natural environments. These aspects are crucial for understanding the ecological relevance of MNP toxicity.

While our meta-analyses found predominantly null effects after limited exposure to virgin MNP in laboratory settings, these results should not be interpreted as evidence that the particles are toxicologically inert. Our study does not encompass environmentally aged plastics, lifelong or multigenerational exposures, or the complex mixtures of sorbed contaminants encountered in real ecosystems. Plastic pollution thus remains a credible ecological and public-health threat, reinforcing the need for regulation and mitigation efforts. Moreover, the generally low methodological and reporting quality of the analyzed studies limits the confidence in current hazard estimates. Well designed and adequately powered experiments are necessary to generate higher quality and more informative data. Further investment is therefore warranted to close critical knowledge gaps.

## 5. Conclusion

In conclusion, while current evidence indicates potential but inconsistent neurotoxic effects of MNPs in fish, future studies with improved reporting, standardized protocols, and broader ecological endpoints are essential for advancing the field and informing environmental regulation.

## Credit authorship contribution statement

**QKZ:** Conceptualization, Methodology, Software, Validation, Formal analysis, Investigation, Data Curation, Writing - Original Draft, Writing - Review & Editing, Visualization; **MGL:** Conceptualization, Methodology, Validation, Investigation, Data Curation, Writing - Original Draft, Writing - Review & Editing, Visualization; **GAM:** Methodology, Investigation; **INL:** Methodology, Visualization; **MEC:** Conceptualization, Methodology, Software, Validation, Formal analysis, Investigation, Data Curation, Writing - Original Draft, Writing - Review & Editing, Visualization, Supervision, Project administration, Funding acquisition; **AP:** Conceptualization, Methodology, Software, Validation, Formal analysis, Investigation, Data Curation, Writing - Original Draft, Writing - Review & Editing, Visualization, Supervision, Project administration; and **APH:** Conceptualization, Methodology, Software, Validation, Formal analysis, Investigation, Data Curation, Writing - Original Draft, Writing - Review & Editing, Visualization, Supervision, Project administration.

## Acknowledgements

The authors are grateful for the financial support by Conselho Nacional de Desenvolvimento Científico e Tecnológico (CNPq) and Coordenação de Aperfeiçoamento de Pessoal de Nível Superior (CAPES).

