## Supplementary material for "Neurological effects induced by micro- and nanoplastics in fish: a systematic review and meta-analysis"

**Table S1.** Stage of development time for *Lates calcarifer, Clarias gariepinus, Danio rerio* and *Sparus aurata*.

| Species | Larva | Fingerling | Juvenile | Adult male | Adult female | Ref |
|---|---|---|---|---|---|---|
| *Lates calcarifer* | 18 hpf | - | 25-30 dph | 3 years | 6-8 years | [34] |
| *Clarias gariepinus* | 60 hpf | 14 dph | | 70 dph | | [35] |
| *Danio rerio* | 72 hpf | 14 dph | 30 dpf | 90 dpf | 90 dpf | [36] |
| *Sparus aurata* | 24 hpf | - | 43-50 dph | 1-2 years | 3 years | [37] |

Lm: (English) Mean length at first maturity

| Study | D1 | D2 | D3 | D4 |
|---|---|---|---|---|
| Ašmonaitė et al., 2020 | + | X | X | X |
| Barboza et al., 2018 | + | X | X | X |
| Barboza, Vieira LR & Guilhermino, 2018 | + | X | X | X |
| Bour et al., 2020 | + | X | X | X |
| Brun et al., 2019 | + | X | X | X |
| Campos et al., 2021 | X | X | X | X |
| Cedervall et al., 2012 | + | X | X | X |
| Chae et al., 2018 | X | X | X | X |
| Chagas et al., 2021 | + | X | X | X |
| Chen et al., 2017 | + | X | X | X |
| Chen et al., 2020 | X | X | X | X |
| Choi et al., 2018 | + | X | X | X |
| Coffin et al., 2020 | X | X | X | X |
| Cormier et al., 2019 | X | X | + | X |
| Cormier et al., 2021 | + | X | + | X |
| Critchell & Hoogenboom, 2018 | + | X | X | X |
| da Costa Araújo, Andrade Vieira & Malafaia, 2020 | X | X | X | X |
| de Oliveira et al., 2021 | + | X | X | X |
| Dimitriadi et al., 2021 | + | X | + | X |
| Ding et al., 2018 | + | X | X | X |
| Ding et al., 2020 | + | X | X | X |
| Estrela et al., 2021 | X | X | X | X |
| Ferreira et al., 2016 | + | X | X | X |
| Fonte, Ferreira & Guilhermino, 2016 | + | X | X | X |
| Guimarães et al., 2021 | + | X | X | X |
| Guven et al., 2018 | + | X | X | X |
| Hu et al., 2021 | + | X | X | X |
| Huang et al., 2021 | + | X | X | X |
| Iheanacho & Odo, 2020 | + | X | X | X |
| Iheanacho et al., 2020 | + | X | X | X |
| Jacob et al., 2019 | + | X | X | X |
| Ji et al., 2020 | + | X | X | X |
| Le Bihanic et al., 2020 | + | X | X | X |
| Lee et al., 2019 | X | X | X | X |
| Limonta et al., 2019 | + | X | + | X |
| Limonta et al., 2021 | X | X | X | X |
| Liu et al., 2019 | X | X | X | X |
| Liu et al., 2021 | + | X | X | X |
| Luís et al., 2015 | X | X | X | X |
| Mak, Ching-Fong Yeung & Chan, 2019 | + | X | X | X |
| Mattsson et al., 2015 | X | X | X | X |
| Mattsson et al., 2017 | X | X | X | X |
| McCormick et al., 2020 | + | X | X | + |
| Miranda, Vieira & Guilhermino, 2019 | X | X | X | X |
| Nanninga et al., 2021 | + | + | + | X |
| Pannetier et al., 2019 | + | X | X | X |
| Pannetier et al., 2020 | + | X | X | X |
| Parenti et al., 2019 | + | X | + | X |
| Parenti et al., 2021 | X | X | X | X |
| Pedersen et al., 2020 | + | X | + | X |
| Pitt et al., 2018a | + | X | X | + |
| Pitt et al., 2018b | + | X | X | X |
| Qiang & Cheng, 2019 | + | X | X | X |
| Qiang & Cheng, 2021 | + | X | X | X |
| Rios-Fuster et al., 2021 | + | X | X | X |
| Rochman et al., 2017 | + | X | X | + |
| Roda et al., 2020 | X | X | X | + |
| Romano et al., 2020 | + | X | X | X |
| Santos et al., 2020 | + | X | + | X |
| Santos et al., 2021a | + | X | X | X |
| Santos et al., 2021b | + | X | X | X |
| Sarasamma et al., 2020 | + | X | X | X |
| Schmieg et al., 2020a | + | X | X | + |
| Schmieg et al., 2020b | X | X | X | X |
| Sheng, Zhang & Zhang, 2021 | + | X | X | X |
| Shi et al., 2021 | + | X | X | X |
| Skjolding et al., 2017 | X | X | X | X |
| Sökmen et al., 2020 | + | X | X | X |
| Trevisan, Uzochukwu & Di Giulio, 2020 | X | X | X | X |
| Umamaheswari et al., 2020 | + | X | X | X |
| Wan et al., 2019 | + | X | X | X |
| Wen et al., 2018 | + | X | X | X |
| Yang et al., 2020a | + | X | X | X |
| Yang et al., 2020b | + | X | X | X |
| Yin et al., 2018 | + | X | X | X |
| Yin et al., 2019 | + | X | X | X |
| Zhang et al., 2020 | + | X | X | X |
| Zhang et al., 2021a | + | X | X | X |
| Zhang et al., 2021b | X | X | X | X |

D1: 1 - Randomization
D2: 2 - Sample size estimation
D3: 3 - Exclusion or inclusion criteria
D4: 4 - Blinding

X No
+ Yes

**Figure S1.** Quality of report assessment of the individual studies.

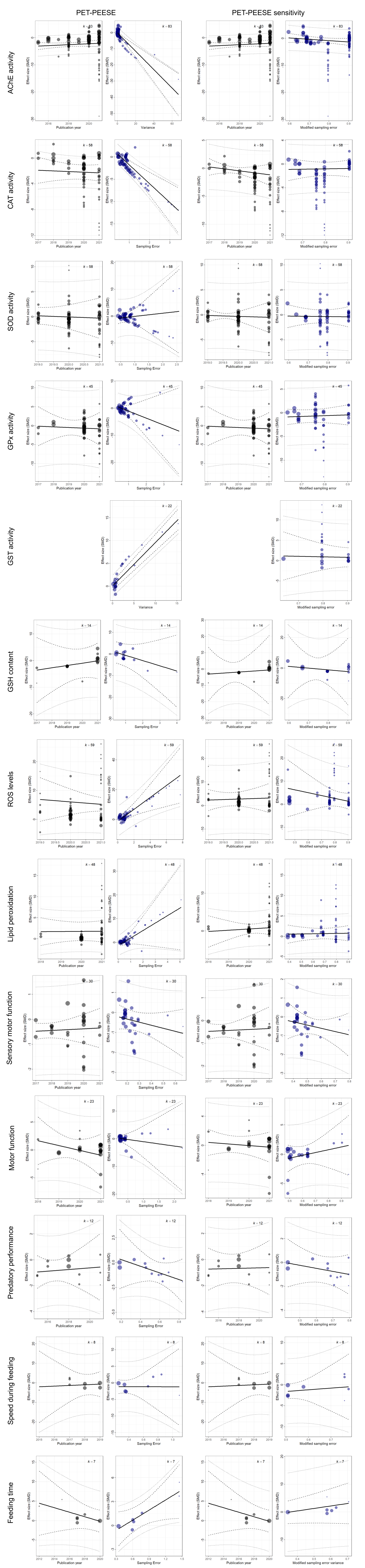


**Figure S2.** Orchard plots of meta-regression with publication bias correction for neurochemical and behaviour outcomes. Gray = small study effects, blue = time-lag bias, filled circle = individual ES scaled by sample size, black line = linear regression with associated 95% confidence interval (dashed line) and prediction intervals (dotted line), k = number of ES

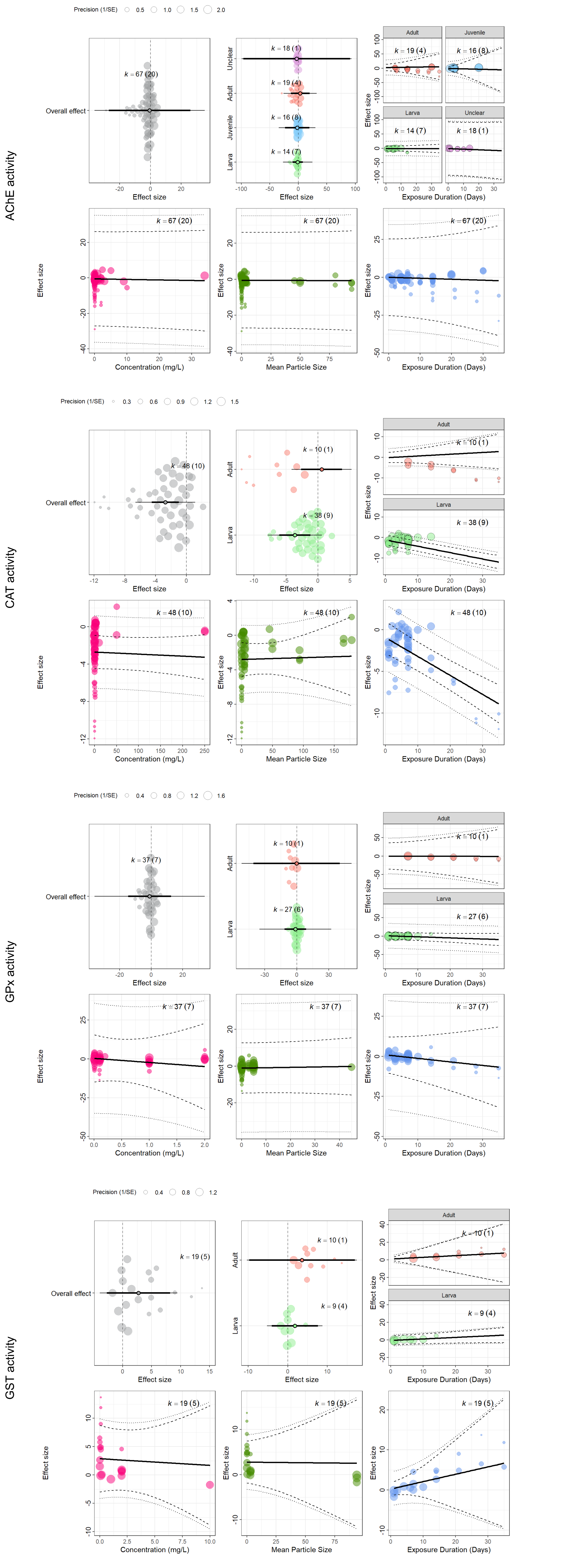


**Figure S3.** Orchard plots of meta-regression with moderators for AChE, CAT and GPx activity.
Superior left and middle plots: filled circle = individual ES scaled by sample size, open circle = point estimate with associated 95% confidence interval (thick black horizontal line) and prediction intervals (thin black horizontal line), k = number of ES (number of reports).
Superior right and inferior plots: black line = linear regression estimate with associated 95% confidence interval (dashed line) and prediction intervals (dotted line), k = number of ES.
